# Native molecular architectures of centrosomes in *C. elegans* embryos

**DOI:** 10.1101/2024.04.03.587742

**Authors:** Fergus Tollervey, Manolo U. Rios, Evgenia Zagoriy, Jeffrey B. Woodruff, Julia Mahamid

## Abstract

Centrosomes organize microtubules that are essential for mitotic divisions in animal cells. They consist of centrioles surrounded by Pericentriolar Material (PCM). Questions related to mechanisms of centriole assembly, PCM organization, and microtubule formation remain unanswered, in part due to limited availability of molecular-resolution structural analyses *in situ*. Here, we use cryo-electron tomography to visualize centrosomes across the cell cycle in cells isolated from *C. elegans* embryos. We describe a pseudo-timeline of centriole assembly and identify distinct structural features including a cartwheel in daughter centrioles, and incomplete microtubule doublets surrounded by a star-shaped density in mother centrioles. We find that centriole and PCM microtubules differ in protofilament number (13 versus 11) indicating distinct nucleation mechanisms. This difference could be explained by atypical γ-tubulin ring complexes with 11-fold symmetry identified at the minus ends of short PCM microtubules. We further characterize a porous and disordered network that forms the interconnected PCM. Thus, our work builds a three-dimensional structural atlas that helps explain how centrosomes assemble, grow, and achieve function.

## Introduction

The centrosome is a conserved microtubule organizing center that is important for building the mitotic spindle. Centrosomes consist of a pair of centrioles, which are cylindrical structures built out of microtubules arranged in a 9-fold symmetry (Boveri, 1887; De Harven and Bernhard, 1956), surrounded by a structurally poorly-defined, micron-scale mass of PCM (Gould and Borisy, 1977).

The composition and molecular architecture of centrioles are well described across many species (LeGuennec et al., 2021; Winey and O’Toole, 2014). The inner centriole is built from a core set of conserved proteins: CPAP/SAS-4, STIL/SAS-5, SAS-6, Plk4/ZYG-1 (Azimzadeh and Marshall, 2010). Surrounding this core are microtubules arranged in either triplets (vertebrates, *Chlamydomonas)* (Chrétien et al., 1997; Kitagawa et al., 2011; Paintrand et al., 1992), doublets (*Drosophila melanogaster* somatic cells) (Gottardo et al., 2015), or singlets (*Caenorhabditis elegans)* (Pelletier et al., 2006; Sugioka et al., 2017). Centrioles duplicate in each cell cycle during S-phase, wherein a procentriole nucleates at an orthogonal orientation from the wall of the mother centriole (Azimzadeh and Bornens, 2007). The procentriole is templated by a structure termed the cartwheel, a central tube from which 9 spokes emanate and nucleate the centriole microtubules. The procentriole matures into the daughter centriole throughout the cell cycle, with the mother and daughter separating to each serve as a new platform for duplication in the newly formed daughter cells. While the molecular factors required for centriole duplication are known, the stepwise structural changes are not defined due to the lack of high-resolution snapshots of intermediates during this process. In the context of the centrosome, the centriole is implicated in imparting mechanical stability to the centrosome (Abal et al., 2005; Le Guennec et al., 2020) and in regulating its replication (Mazia et al., 1960; Nigg and Holland, 2018; Sluder and Begg, 1985).

The PCM occupies the bulk of the centrosome and is responsible for the organization and assembly of microtubules (Gould and Borisy, 1977). In interphase, the PCM consists of thin patterned layers only a few hundred nanometers in diameter (Fu and Glover, 2012; Lawo et al., 2012; Mennella et al., 2012; Sonnen et al., 2012). In mitosis, the PCM increases in size and “matures” by accumulating components needed for microtubule nucleation and anchoring, expanding its capacity for microtubule formation to build the mitotic spindle (Conduit et al., 2014; Decker et al., 2011; Khodjakov and Rieder, 1999). One such component is the γ-tubulin ring complex (γ-TuRC), which is important for templating the growth of microtubules, stabilizing their minus ends (Moritz et al., 2000, 1995), and helping to organize the entire spindle (O’Toole et al., 2012). Other components include TPX2, ch-TOG, and CLASP family proteins that assist with microtubule growth, stabilization, and maintenance (Tsuchiya and Goshima, 2021).

The PCM is enriched in long coiled-coil proteins, such as SPD-5 in *C. elegans* (Hamill et al., 2002; Woodruff et al., 2017, 2015), Cnn (centrosomin) in *Drosophila* (Timothy et al., 1999), and CDK5RAP2 in humans (Fong et al., 2007). SPD-5 and Cnn are sufficient to multimerize into micron-scale structures *in vitro*, suggesting that the PCM is built through self-assembly of such scaffolding proteins (Feng et al., 2017; Woodruff et al., 2015). This process is potentiated by Polo family kinases, SPD-2/Cep192, and Aurora A kinase (Conduit et al., 2014; Decker et al., 2011; Haren et al., 2009; Joukov et al., 2014; Woodruff et al., 2015). Despite having a comprehensive parts list for the PCM, how these proteins are organized within the expanded mitotic PCM remains a mystery. Although much work has been carried out using room temperature electron microscopy (EM) on the centrosome in a number of organisms, the PCM typically appears as an unstructured volume surrounding the centrioles (Bowler et al., 2019; De Harven and Bernhard, 1956; Debec et al., 1999; O’Toole et al., 2012; Pelletier et al., 2006, 2004; Redemann et al., 2017; Sugioka et al., 2017). A notable exception to this was found in salt-stripped centrosomes of the surf clam *Spisula solidissima*, in which a network of 12-15 nm long fibers form an underlying scaffold that was termed the ‘centromatrix’ (Schnackenberg et al., 1998). It however remains unclear whether this appearance is an artefact of sample preparation or is a species-specific phenomenon. Thus, it is unknown whether the centromatrix exists under native conditions and whether it may constitute a general feature of the PCM across organisms.

*C. elegans* has served as a powerful model for the study of centrosomes, especially during embryonic development, owing to its relative compositional simplicity compared to that in other organisms, such as *Drosophila melanogaster* or humans (reviewed in (Lattao et al., 2017) and (Vasquez-Limeta and Loncarek, 2021) respectively). As such, some of our best understanding of centriole development and PCM function arises from this nematode. In particular, the process of centriole biogenesis is well documented and requires 6 proteins acting in a specific order. First, SAS-7 (Sugioka et al., 2017) and SPD-2 (Pelletier et al., 2004) are recruited to the mother centriole. These proteins recruit the kinase ZYG-1 (Plk4 homolog) which sets the site of procentriole formation by recruiting SAS-5, SAS-6, and then SAS-4 (Dammermann et al., 2004; Pelletier et al., 2006). SAS-6 and SAS-5 are of particular importance in templating the 9-fold symmetry of the growing centriolar microtubules. SAS-6 was originally suggested to form a helical assembly in the center of the procentriole (Hilbert et al., 2013). However, later super-resolution microscopy work showed that the N-terminal heads of SAS-6, in complex with SAS-5, form a tube structure (Woglar et al., 2022). For the procentriole to mature, PCMD-1, SAS-1, SAS-2, and SPD-5 must also be recruited. Expansion microscopy of *C. elegans* gonads revealed that SPD-2, SPD-5, and PCMD-1 localize adjacent to the microtubules in the centriole (Woglar et al., 2022). This and previous studies using room temperature EM defined a centriolar ‘paddlewheel’, a region of poorly defined electron dense material, budding off from the a-tubule and spreading the entire length of the centriole (Dammermann et al., 2004; Pelletier et al., 2006; Sugioka et al., 2017; Woglar et al., 2022). The paddlewheel is the only known chiral structure in the embryonic *C. elegans* centriole, yet its identity and fine structure remain unknown.

As for the PCM, the relatively small number of proteins involved in *C. elegans* culminated in the reconstitution of an *in vitro* minimal system capable of nucleating microtubule asters (Woodruff et al., 2017). PCM maturation in *C. elegans* requires SPD-2 (Decker et al., 2011; Kemp et al., 2004; Pelletier et al., 2004), and the kinase PLK-1 (Ohta et al., 2021), which recruit SPD-5 proteins and enhance their ability to self-assemble into a supramolecular scaffold (Wueseke et al., 2016). Once expanded, the PCM scaffold recruits client proteins such as γ-tubulin (Kollman et al., 2015; Moritz et al., 1995), TPXL-1 (Özlü et al., 2005), and the ch-TOG family ZYG-9 (Matthews et al., 1998), that concentrate tubulin (Baumgart et al., 2019; Woodruff et al., 2017) and nucleate microtubules to help form the mitotic spindle. Upon the completion of mitosis, the expanded PCM disassembles due to phosphatase-mediated material weakening of the scaffold and force-driven dispersal (Enos et al., 2018; Mittasch et al., 2020). Open questions remain as to the underlying architecture of the PCM scaffold and how it may change in adaptation to the different PCM functions during its expansion, maturation, and disassembly.

To provide insight into the structural evolution of centrosomes throughout the cell cycle, here we use cryogenic electron tomography (cryo-ET) on dividing *C. elegans* embryonic cells. In contrast to traditional EM that was employed in most ultrastructural studies of centrosomes to date, this method allows the preservation of fine detail for cellular structures visualized in close-to-native state (Adrian et al., 1984). We thereby characterize changes to the centrosome architecture, visualize the centriole structure at different stages of maturation, investigate the architectural role of the centrosome in organizing microtubules, and identify the *C. elegans* γ-tubulin ring complex in the PCM. Finally, we resolve a PCM-localized matrix that comprises a disorganized meshwork with pore sizes capable of accommodating PCM client proteins.

## Results

### A Cryo-ET pipeline to visualize centrosomes in *C. elegans* embryonic cells

Structural studies of cells by cryo-ET require vitrification to preserve their native state, followed by thinning to approximately 200 nm using cryo-focused ion beam (cryo-FIB) (Schaffer et al., 2017) to enable molecular-resolution imaging in a transmission electron microscope (TEM). Cryo-fluorescence imaging can be used to guide the milling process and/or cryo-ET imaging to target specific regions of interest (Arnold et al., 2016; Klumpe et al., 2021). However, sample thickness is a fundamental consideration: samples must be thinner than 10 μm for effective vitrification by routine plunge-freezing and cryo-FIB lamellae preparations, which presents a limitation for *C. elegans* worms and embryos (up to 100 μm and 40 μm thickness, respectively). Although high pressure freezing can vitrify samples in this thickness range, the preparation of cryo-FIB lamellae from voluminous specimen is yet to be fully streamlined (Mahamid et al., 2015; Schaffer et al., 2019; Schiøtz et al., 2023).

Therefore, to visualize centrosomes in their native state and to avoid the limitations of freezing large, intact *C. elegans* embryos, we adapted a protocol for isolation of cells from early-stage embryos (Material and Methods, Supplementary Figure 1A) (Christensen et al., 2002). The dissociated cells were viable in culture and continued to divide for at least 160 minutes (Supplementary Figure 1B). The cells were deposited on EM grids, plunge frozen, thinned by cryo-FIB milling and imaged by cryo-ET. Using untargeted cryo-FIB milling, we obtained 5 tomograms of interphase centrosomes, identified based on the stereotypical structural signature of the centrioles and the presence of an intact nuclear envelope (Figure 1A, Supplementary Figure 2A). To target mitotic centrosomes, the plunge-frozen grids were imaged using a cryo-confocal fluorescence microscope (Supplementary Figure 1C). Using a *C. elegans* line expressing GFP::SPD-5 (PCM marker) and H2B::mCherry (DNA marker), we identified mitotic cells and the relative positioning of the centrosome and mitotic chromosomes to aid targeting of the cryo-FIB milling process (Supplementary Figure 1D). The cell cycle stages were also determined using a combination of the cryo-fluorescence data and structural information from the cryo-TEM images (e.g., how intact the nuclear envelope appeared). Overall, our approach allowed us to obtain 12 tomograms of centrosomes in interphase (5), prophase (1), metaphase (3), anaphase (2), and telophase (1) (Table 1). We used these tomograms to characterize the evolution of the molecular architectures of centrosomes across the cell cycle.

**Figure 1.**
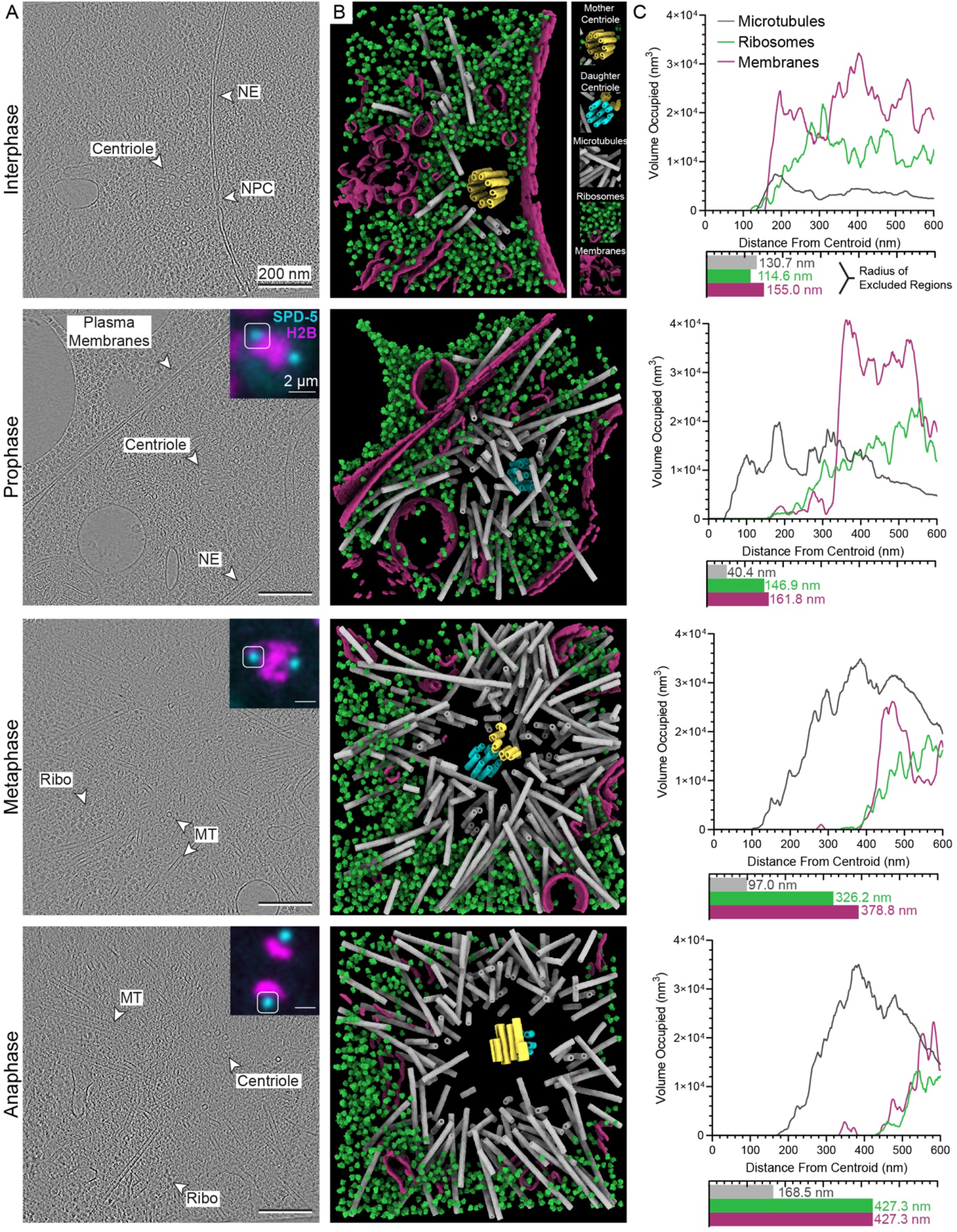
Cryo-electron tomograms reveal native centrosome architectures across the cell cycle in *C. elegans* embryonic cells. Cell cycle stage is indicated on the left. A) Tomographic slices. Features of interest are labelled with arrowheads. NE: Nuclear Envelope, NPC: Nuclear Pore Complex, Ribo: Ribosomes, MT: Microtubules. Insets show correlated fluorescence images (maximum intensity projection of the cryo-confocal stack; GFP::SPD-5: cyan; H2B::mCherry: magenta) at the same orientation as the tomographic data. B) 3D annotation of the data shown in A. Mother centriole microtubules: yellow, daughter centrioles: cyan (note central tube depicts the cartwheel), microtubules: grey, ribosomes: green, and membranes: purple. C) Integrated volumes of microtubules, ribosomes, and membranes as a function of distance from their corresponding centroids. The exclusion zones, defined as the last distance at which each of the features is not detected, is recorded below. The remaining tomograms and annotations analyzed in this work are provided in Supplementary Figure 2.

**Table 1.**
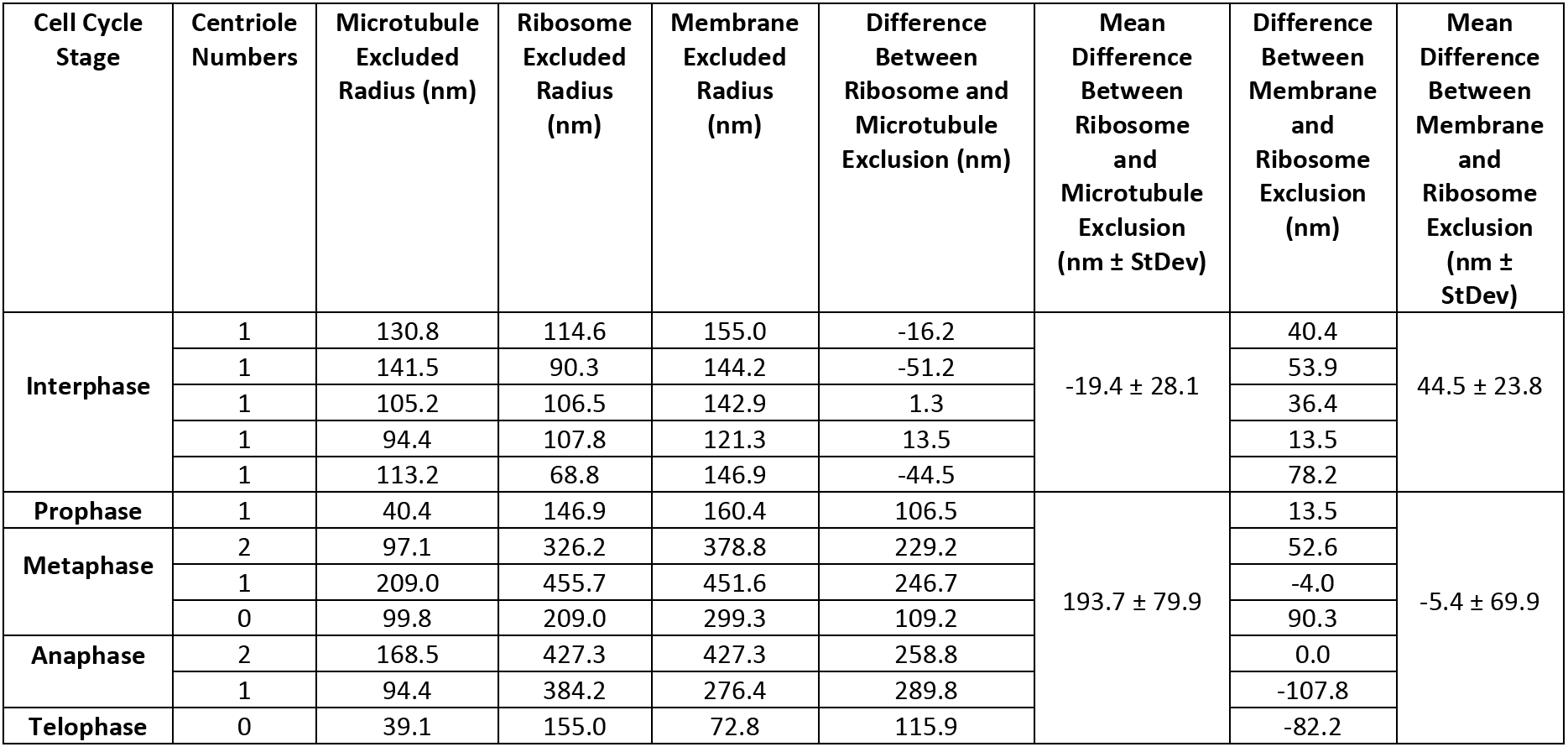
Overview of centrosomes and structural zones analyzed in this study.

### Centrosome architecture at molecular resolution

We identified centrosomes in our tomograms based on the presence of centrioles, and in the case of mitotic centrosomes, through the registration of the cryo-fluorescence data (Figure 1A, Supplementary Figure 1D, Supplementary Figure 2). Since the *C. elegans* centriole is reported to be roughly 150 nm in length and 100 nm in diameter (Dammermann et al., 2004; Pelletier et al., 2006; Sugioka et al., 2017; Woglar et al., 2022), retention of a complete centriole pair within the roughly 200 nm thick lamella is unlikely. Nevertheless, partial centrioles, single complete centrioles, or partial centriole pairs were visible (Table 1). To aid quantitative analysis of the centrosome arrangement, we performed segmentation of different cellular features including centrioles, microtubules, membranes, and ribosomes (Figure 1B), using a combination of manual, semi-automated correlation-based segmentation, and supervised deep-learning based annotation (Material and Methods).

Our data revealed that centrosome architectures can be defined based on concentric ‘zones’. At the center of each centrosome was a centriole surrounded by an area devoid of microtubules and ribosomes (“microtubule-excluded zone”) and an area rich in centrosomal microtubules but without ribosomes (“ribosome-excluded zone”). As ribosomes are abundant in the cytoplasm and not recognized as PCM components, we hypothesize that the edge of the ribosome-excluded zone represents a minimum estimate of the outer boundary of PCM. We observed membranes near the interface between PCM and the cytoplasm. Here, we refer to the area surrounding a centriole that lacks membranes as the “membrane-excluded zone”.

We then examined how these zones change throughout the cell cycle. We calculated the radii of the microtubule, ribosome, and membrane-excluded zones based on the segmentation, which were defined as the last voxel that does not contain the cellular structure of interest from the centroid defined for each of the structures (Figure 1C, Material and Methods). In interphase cells, where the centrosome center was defined as the center of the centriole, the ribosome-excluded zone radii ranged from 68.8 nm and 114.6 nm. The microtubule-excluded radius was, on average 19.4 ± 28.1 nm larger than the ribosome-excluded radius (Figure 1C; Table 1). We conclude that microtubules are primarily nucleated at the surface of interphase PCM, in agreement with previous room-temperature EM studies (O’Toole et al., 2012; Pelletier et al., 2004; Redemann et al., 2017). The membrane-excluded region was larger than both the microtubule- and ribosome-excluded areas. All interphase centrosomes were adjacent to a nuclear pore embedded within the nuclear envelope, separated by a mean distance 191.5 ± 5.0 nm from the center of the centriole (Supplementary Figure 2A). This finding supports previous studies showing that PCM is physically linked to the nucleus through dynein motors anchored to the nuclear pore complex (Guo and Yixian, 2015).

In mitotic centrosomes, the ribosome-excluded radii ranged from 146.9 nm (prophase) to 455.7 nm (metaphase), indicating an increase in PCM size compared to interphase. In contrast to interphase centrosomes, the microtubule-excluded zone was more easily distinguished from the ribosome-excluded zone; the mean difference between the microtubule- and ribosome-excluded radii was 193.7 ± 79.9 nm (Table 1, Figure 1C). However, the microtubule-excluded zone was on average 70% smaller in prophase compared to interphase (Table 1, Figure 1C), suggesting that maturing centrosomes can nucleate microtubules within the PCM, not only at the surface. The microtubule-excluded zone increased in size as cells progressed into anaphase, which could be due to an increase in microtubule-mediated pulling forces that are known to physically stretch PCM (Enos et al., 2018; Megraw et al., 2002). We calculated a mean difference of only −5.4 ± 69.9 nm between the radii of the ribosome-excluded region and the membrane-excluded region (Table 1). Thus, the zone of membrane exclusion is nearly equivalent to that of ribosome exclusion in mitosis. Previous reports suggested that mitotic centrosomes are surrounded by an extension of the endoplasmic reticulum termed the ‘centriculum’ (Maheshwari et al., 2023). However, these membranes do not fully enclose the PCM in our data, indicating that the outer boundary of PCM is not entirely defined by membranes; thus, centrosomes should not be considered membrane-enclosed organelles. We conclude that the PCM excludes both ribosomes and membranes.

In summary, our cryo-ET approach identified architectural zones that change in size during centrosome maturation. The presence of concentric microtubule- and ribosome-excluded zones within the centrosome is consistent with previous traditional room-temperature EM studies of intact *C. elegans* embryos (O’Toole et al., 2012; Pelletier et al., 2004; Redemann et al., 2017), supporting the utility of studying dissociated embryonic cells. We thus proceeded to analyze native centriole and PCM structures at high resolution.

### *In situ* analysis of the developing centriole

The *C. elegans* centriole has been thoroughly investigated by room temperature EM (Dammermann et al., 2004; Pelletier et al., 2006, 2004; Sugioka et al., 2017; Woglar et al., 2022), providing a rich source of data on the ultrastructure of this molecular assembly. However, the associated sample preparation involving chemical fixation, dehydration, staining, and resin embedding may cause alterations that obscure their fine structural features (Kellenberger et al., 1992; Murk et al., 2003).

In cryo-tomograms where two centrioles could be seen simultaneously in a single centrosome, we found one centriole to be positioned 90° relative to the long axis of the other. We assigned the mother and daughter centrioles based on this relative orientation, with the daughter centriole positioned perpendicular to the cylinder wall of the mother (Figure 2A). The daughter centrioles exhibited features similar to those described in the literature, including the presence of singlet microtubules and an inner tube structure (Figure 2B, Supplementary Figure 3). The size and shape of this tube are consistent with previous *in vitro* studies of SAS-6 in *C. reinhardtii*, (Hatzopoulos et al., 2021), and of a complex of SAS-5/SAS-6 N terminal heads in *C. elegans* (Woglar et al., 2022). For consistency of terminology with other organisms, we thus refer to this structure as the “cartwheel” (Woglar et al., 2022), as opposed to the outdated term “inner tube” (O’Toole et al., 2012; Pelletier et al., 2006; Sugioka et al., 2017). We observed a central tube in both the mother and daughter centrioles, with detectable projections emanating towards the centriole microtubules most prominently in mother centrioles (tube spikes, Figure 2B, Supplementary Figure 3). The mother centriole exhibited several previously unknown features; instead of the paddlewheel, a proteinaceous density reported in room temperature EM (Pelletier et al., 2006; Sugioka et al., 2017), we observed a partial b-tubule. Furthermore, a density around the centriole circumference formed a star shape. This star density was most distinguishable at the junction between the a- and b-tubules (a structure we call the ‘star joint’) and at the tip of the b-tubule (‘star tip’).

**Figure 2.**
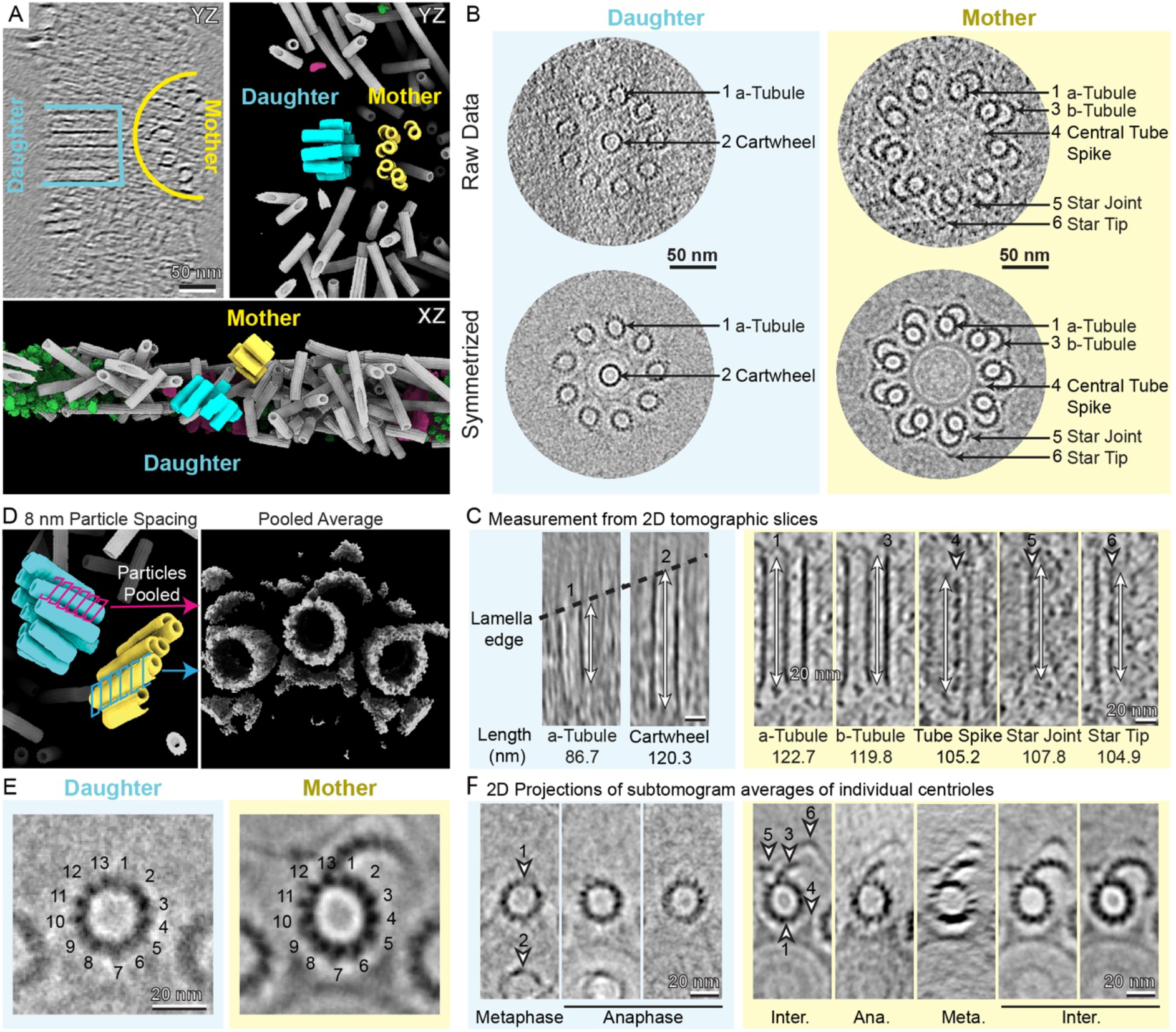
Native centriole structures at different stages of maturation. A) Segmentations of the tomograms allow identification of the mother (yellow) and daughter (cyan) centrioles. The daughter centriole is positioned at a 90° angle off the side of the mother centriole in both the XY and YZ planes. Color scheme as in Figure 1B. B) Cross section views of the mother and the daughter centrioles, both unsymmetrized and with 9-fold symmetry applied. The complete set of centrioles analyzed in this work is provided in Supplementary Figure 3A. Arrows indicate key features: the a-tubule (1), cartwheel (2), b-tubule (3), central tube spikes (4), outer star joint (5), and outer star tip (6). C) Lengths of centriole structural features as determined through intensity line scans in the raw tomograms. Distances indicated are from the averages detailed in Table 2. D) Subtomogram averaging of centrioles. Subtomograms were sampled every 8 nm along each centriole a-tubule, pooled together and aligned into a single average. Averages were then split into pooled daughter and mother centrioles (E) or for each centriole (f). E) 2D projections of pooled daughter and mother subtomogram averages. Indicated are protofilament numbers in the a-tubule. F) Gallery of 2D projections of per-centriole subtomogram averages, corresponding to the features detailed in B and C. Cell cycle stages are indicated on the bottom.

We next quantified the sizes of the newly-identified structures by measuring intensities using line scans along the centrioles in the tomograms (Materials and Methods, Figure 2C, Table 2). In the daughter centrioles, the cartwheel averaged 120.3 nm in length, extending on average 33.6 nm beyond the a-tubules toward the mother centriole. We did not visualize complete daughter centrioles in this study (that are not cut off by the lamella edge), and therefore the total length of the a-tubules could not be determined (measured lengths were at least 86.7 nm). Analysis of the mother centriole revealed a total length of 122.7 nm for the a-tubule and 119.8 nm for the b-tubule. The spikes of the central tube spanned 105.2 nm of the centriole length and exhibited repeats of variable distance. The star joint covered 107.8 nm, and the tip 104.9 nm of the centriole length, both exhibiting interruptions in their density along their length. Each of the features unique to the mother (b-tubule, tube spikes, star joint, and star tip) were found at an equal distance from the proximal and distal ends of the centriole and therefore appear to lack polarity.

**Table 2.**
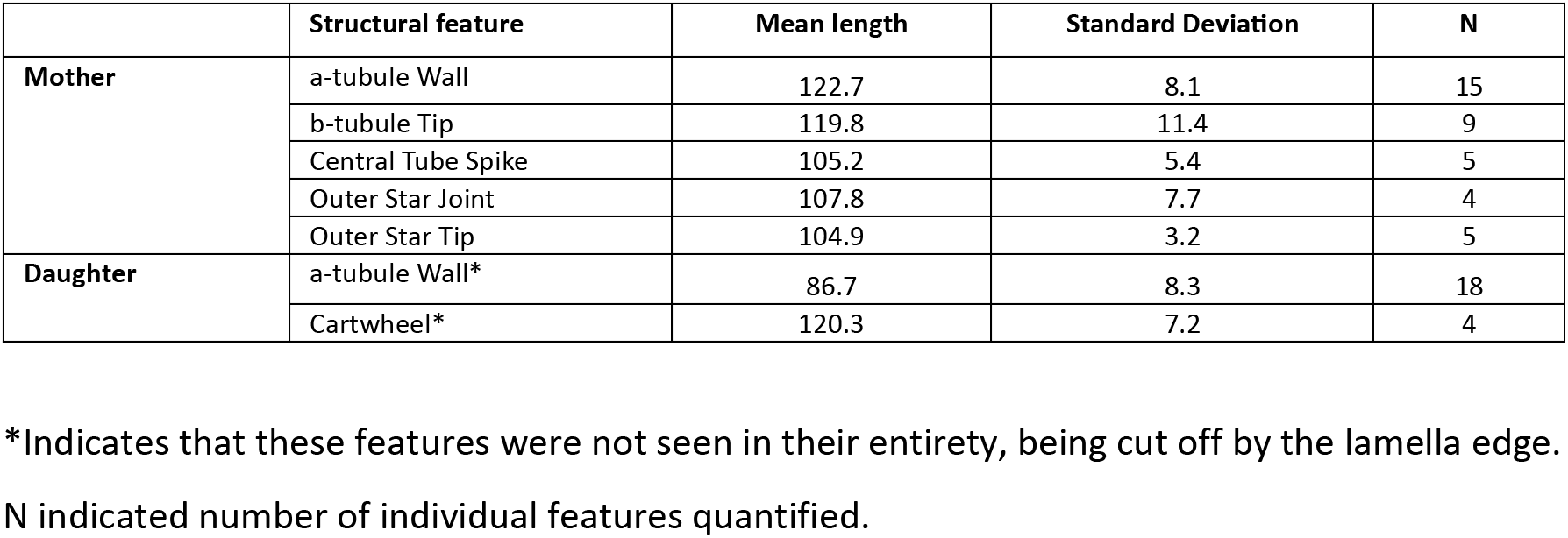
Quantification of centriole structural features analyzed in this study.

To provide higher resolution structural detail, we next performed subtomogram averaging by picking a total of 800 particles with 8 nm spacing along each a-tubule in all 12 centrioles (Materials and Methods, Figure 2D). Due to the low particle numbers arising from the short lengths of the centrioles, high resolution averages were not obtained, with the map achieving a final resolution of 44 Å (Supplementary Figure 3B). Nevertheless, 2D projections of per-centriole, non-symmetrized averages confirmed the presence of microtubule doublets and the outer star in mother centrioles only (Figure 2E, F). Additionally, a-tubules consisted of 13 protofilaments in both mother and daughter centrioles, which contrasts with the well-described prominence of 11 protofilament microtubules in *C. elegans* (discussed in detail in the next subsection (Figure 2E) (Chaaban et al., 2018; Chalfie and Thomson, 1982). b-tubules in the mother centrioles contained 3-9 protofilaments, with no apparent correlation between protofilament number and cell cycle stage. In no case did the b-tubule fully close onto the a-tubule (Figure 2F). We conclude that maturing centrioles in *C. elegans* appear to lose the cartwheel structure, while they build a central tube, an incomplete b-tubule and an outer star structure superficial to the b-tubule.

### Organization of centrosome microtubules and γ-tubulin ring complexes

The mitotic PCM nucleates and organizes thousands of microtubules to build a functional spindle (Redemann et al., 2017). Our 3D segmentations of centrosome from the cryo-ET data enabled us to investigate, in high detail, the local organization of these microtubules. We first investigated the homogeneity in microtubule protofilament number, previously reported to be 11 in *C. elegans* from room-temperature EM studies (Chaaban et al., 2018; Chalfie and Thomson, 1982). We sampled 8 nm segments along the long axis of the microtubules, aligned and averaged a total of 19,339 subtomograms into a single map with 26 Å resolution (Materials and Methods, Figure 3A, Supplementary Figure 4A, B). The data was then split into 830 per-microtubule maps (Figure 3A), and 2D projections of the averages were compared to references of microtubules of 11 to 15 protofilaments assembled *in vitro* (Sui and Downing, 2010)(Figure 3B). Of all the microtubules analyzed (mean length = 149 ± 109 nm), all except 2 consisted of 11 protofilaments (Table 3). Thus, the number of microtubule protofilaments is highly consistent in the *C. elegans* PCM. Microtubule polarity, as determined by examining the protofilament skew (Sosa and Chrétien, 1998) (Figure 3C), was similarly conserved, with 80% of microtubules oriented with their growing plus ends away from the centrosome centroid (plus-end out, Table 3), in agreement with previous reports by room temperature EM (Redemann et al., 2017). The minor fraction of microtubules with the opposite polarity (minus-end out) were significantly shorter than plus-end out microtubules (mean length = 124.5 ± 10.5 nm versus 217.8 ± 110.9 nm; p = 2.62*10^-23^, 1-tailed t test) and rarely extended beyond the dense region of microtubules between the ribosome and microtubule excluded zones of the centrosome (Supplementary Figure 4C).

**Figure 3.**
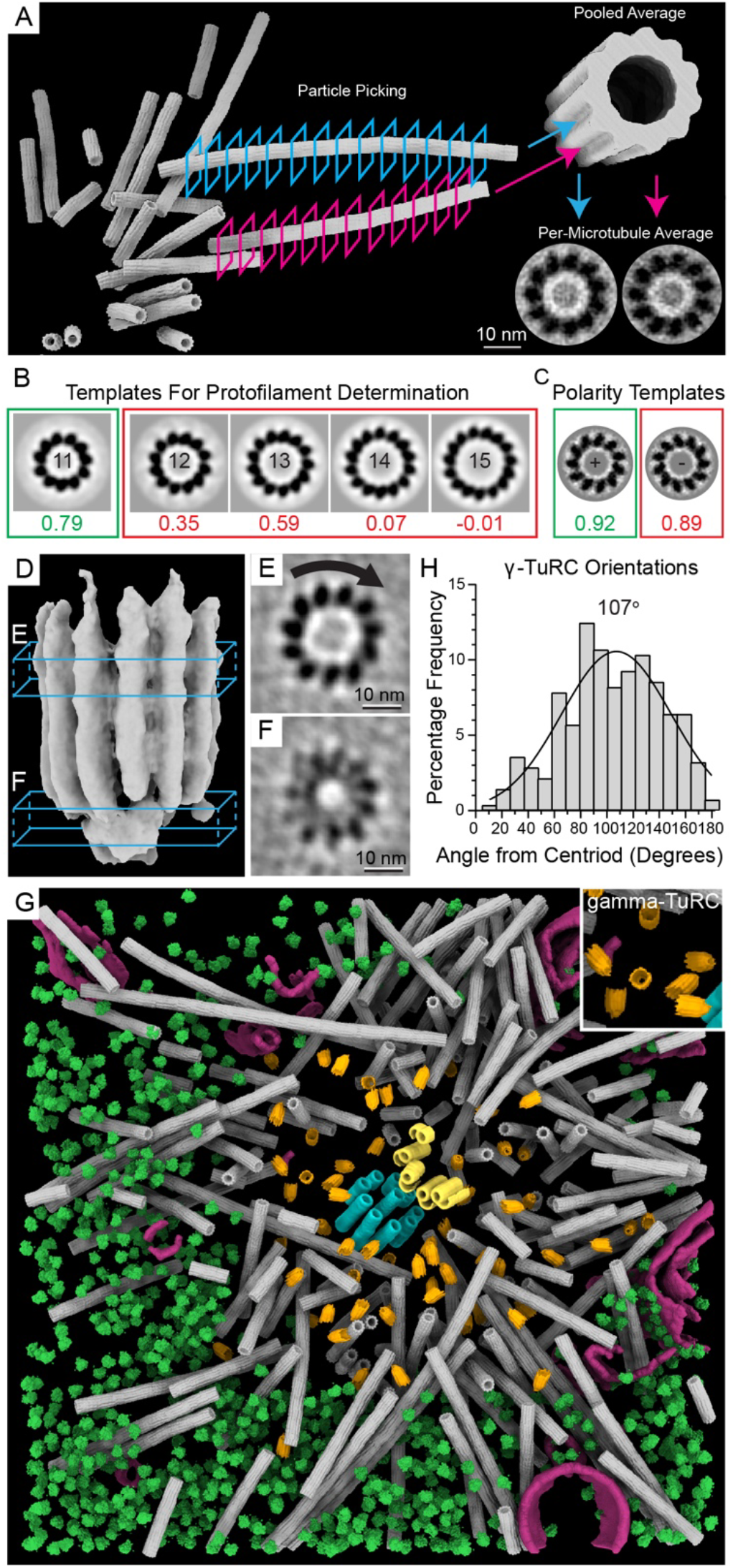
Atypical γ-TuRCs cap and organize 11-protofilament microtubules within the PCM. A) Schematic of the microtubule averaging workflow: microtubules were sampled at 8 nm spacing along each filament to generate subtomograms, which were pooled into a single average. Once aligned, these were separated back into per-microtubule averages and projected along their long axis to increase the signal-to-noise ratio in their cross-sectional view. B) Each per-microtubule average was aligned and cross-correlated with templates corresponding to 11, 12, 13, 14, and 15 protofilament microtubule templates (from EMD-5191 to EMD-5195, (Sui and Downing, 2010)), and the result with the highest cross correlation score was recorded as the protofilament number. C) Each per-microtubule average was aligned and cross-correlated with templates corresponding to positive and negative 11-protofilament microtubule skews. The highest cross correlation score was recorded as the polarity. D) Subtomogram average of the γ-TuRCs, with a capping region at the bottom and a wider microtubule bound section at the top. Boxed regions shown in cross sections in E and F. E) 2D projection of the microtubule bound region of the average, with the skew indicating positive polarity, meaning that the minus end is stabilized by the capped region. F) 2D projection of the lower capped region of the average, showing a ring structure composed of 11 subunits. G) γ-TuRC averages mapped back into the data showed no preferred localization in the centrosome. Note that only γ-TuRC at ends of short microtubules were detected, processed, and displayed. The remaining tomograms and annotations analyzed in this work are provided in Supplementary Figure 4. H) Orientations of γ-TuRCs were determined through the cross product of the vectors along the long axis of the γ-TuRC and the ribosome excluded centroid. Gaussian fitting was applied and the mean value indicated.

**Table 3.** Summary of microtubule protofilament numbers and polarities seen in this study.

| Cell Cycle Stage | Total microtubule number | Number of microtubules with 11 Protofilaments | Number of microtubules with plus-end out orientation | Percentage of microtubules with plus-end out orientation |
| --- | --- | --- | --- | --- |
| Prophase | 72 | 72 | 50 | 69% |
| Metaphase | 169 | 168 | 145 | 86% |
|  | 49 | 49 | 38 | 76% |
|  | 162 | 161 | 114 | 70% |
| Anaphase | 132 | 132 | 100 | 76% |
|  | 196 | 196 | 183 | 93% |
| Telophase | 50 | 50 | 37 | 74% |
| All | 830 | 828 | 755 | 80% |

Our data show that centriole and PCM-nucleated microtubules differ in protofilament numbers (13 versus 11, respectively). We thus asked how is microtubule protofilament number controlled within cellular sub compartments that reside within small distances. Previous work has shown that the majority of microtubules assembled *in vitro* from purified *C. elegans* tubulin have 12-13 protofilaments (Chaaban et al., 2018). Thus, the intrinsic properties of α/β tubulin could explain the geometry of the centriole microtubules but not PCM microtubules. We hypothesized that the 11-protofilament configuration is determined by a PCM-localized nucleating complex with a matching geometry. In most animal cells, nucleation of PCM-based microtubules is dependent on the γ-tubulin ring complex (γ-TuRC). Key members of this complex (e.g. γ-tubulin, MZT-1, GIP-1, GIP-2) localize to *C. elegans* PCM (Ohta et al., 2021; Sallee et al., 2018), yet a complete *C. elegans* γ-TuRC structure has not been reported. We observed numerous short microtubule segments (< 40 nm) capped with a cone-shaped structure within the microtubule-excluded region of centrosomes. These short capped microtubule segments provided an opportunity for performing structural analysis of putative γ-TuRCs, as attempts to localize the complex using template matching with a reference of the human γ-TuRC structures (Consolati et al., 2020; Liu et al., 2020; Wieczorek et al., 2020; Würtz et al., 2022; Zimmermann et al., 2020) were unsuccessful. We thus generated an average from 281 particles, which represent the γ-TuRC complexes that could be manually localized in the data due to their association with short microtubule ends (Figure 3D). We note that free γ-TuRCs were not detected in the data, possibly due to technical limitations in template matching, and that caps of longer astral microtubules could not be aligned in our subtomogram averaging procedure, likely due to the strong density from the microtubule.

The 38 Å resolution average (Supplementary Figure 4D, E) showed a 30 nm long, 11-protofilament microtubule segment connected to a circular cap region. Attempts to identify caps not connected to these very short microtubule segments through classification were unsuccessful, likely due to the small particle number. The protofilament skew on the short microtubule segment revealed the minus end to be located at the circular cap (Figure 3E) (Sosa and Chrétien, 1998), with the cap region containing a prominent hole at the end (Figure 3F, Supplementary Figure 4E). We then compared these caps with structures of the human y-TuRC, which exhibits 13-fold symmetry (Consolati et al., 2020; Liu et al., 2020; Wieczorek et al., 2020; Würtz et al., 2022; Zimmermann et al., 2020). The cone-shaped tip of human γ-TuRC (Wieczorek et al., 2020) (PDB 6V6S) consisting of the N-terminal domains of the GCP subunits, fitted well within our cap average (Supplementary Figure 4F). However, locations distal to the tip fitted poorly. This is possibly due to differences between microtubule protofilament number in humans (commonly 13) and *C. elegans* (11) (Chaaban et al., 2018; Chalfie and Thomson, 1982), or could be attributed to the flared conformation observed in the human y-TuRC complexes derived from reconstituted material in the absence of microtubules (Liu et al., 2020; Wieczorek et al., 2020). Thus, our results reveal that *C. elegans* build γ-TuRCs that match the 11-fold symmetry of PCM microtubules. Importantly, our average represents an active conformation of the γ-TuRC, as indicated by the bound microtubules.

How are the γ-TuRCs oriented with respect to the centrosome? Projected back into the data, these complexes were randomly positioned throughout the ribosome-excluded zone (Figure 3G, Supplementary Figure 4G). This observation does not agree with the notion that the centrosome is delineated by γ-TuRCs as has been previously suggested (O’Toole et al., 2012), but supports the notion that γ-TuRCs are positioned throughout the volume of the centrosome (Moritz et al., 1995). To quantify γ-TuRC orientations with respect to the centrosome architecture, we measured the angle between the vector pointing for the centroid of the ribosome excluded zone to the center of each γ-TuRC, and the vector along the short microtubule segment. The resulting histogram is shown in Figure 3H with a mean angle of 107.2° ± 41.8°. This shows a wide distribution of angles, with a skew towards higher numbers, indicating a slight preference for γ-TuRCs to be oriented away from the centroid, but likely insufficient to explain the large preference for the plus-end out orientation we measured PCM-associated microtubules (Table 3).

We observed no centrioles with capped microtubule ends, suggesting an absence of γ-TuRC. In contrast, all detected γ-TuRCs were bound to a microtubule with 11 protofilaments. We conclude that *C. elegans* embryos build atypical γ-TuRCs with 11-fold symmetry, and we propose that this symmetry then sets protofilament number in non-centriolar microtubules.

### Identification and properties of a porous interconnected centrosome matrix

*C. elegans* PCM forms through self-assembly of the coiled-coil protein SPD-5 (Hamill et al., 2002; Woodruff et al., 2017, 2015). Yet, the meso-scale architecture of the functional PCM underpinned by long-range SPD-5 interactions remains largely unresolved. Here, we use our cryo-ET data to characterize this assembly.

Within the PCM of both interphase and mitotic centrosomes, we identified thin filamentous structures throughout the ribosome-excluded zones (Figure 4A, Supplementary Figure 5). These differed from known components such as microtubule segments. To characterize these filaments, we first masked all previously identified large components (centrioles, microtubules, ribosomes, membranes) out of the tomographic volume to more accurately trace the fine filaments without erroneously tracing these strongly contrasting features (Materials and Methods). Only tomograms in which these filaments were most visually distinct, due to suitably thin lamella and good contrast, were selected for analysis; in total, 7 tomograms were segmented: 3 in interphase, 2 in metaphase, and 2 in anaphase (Figure 4B, Supplementary Figure 5A, B).

**Figure 4.**
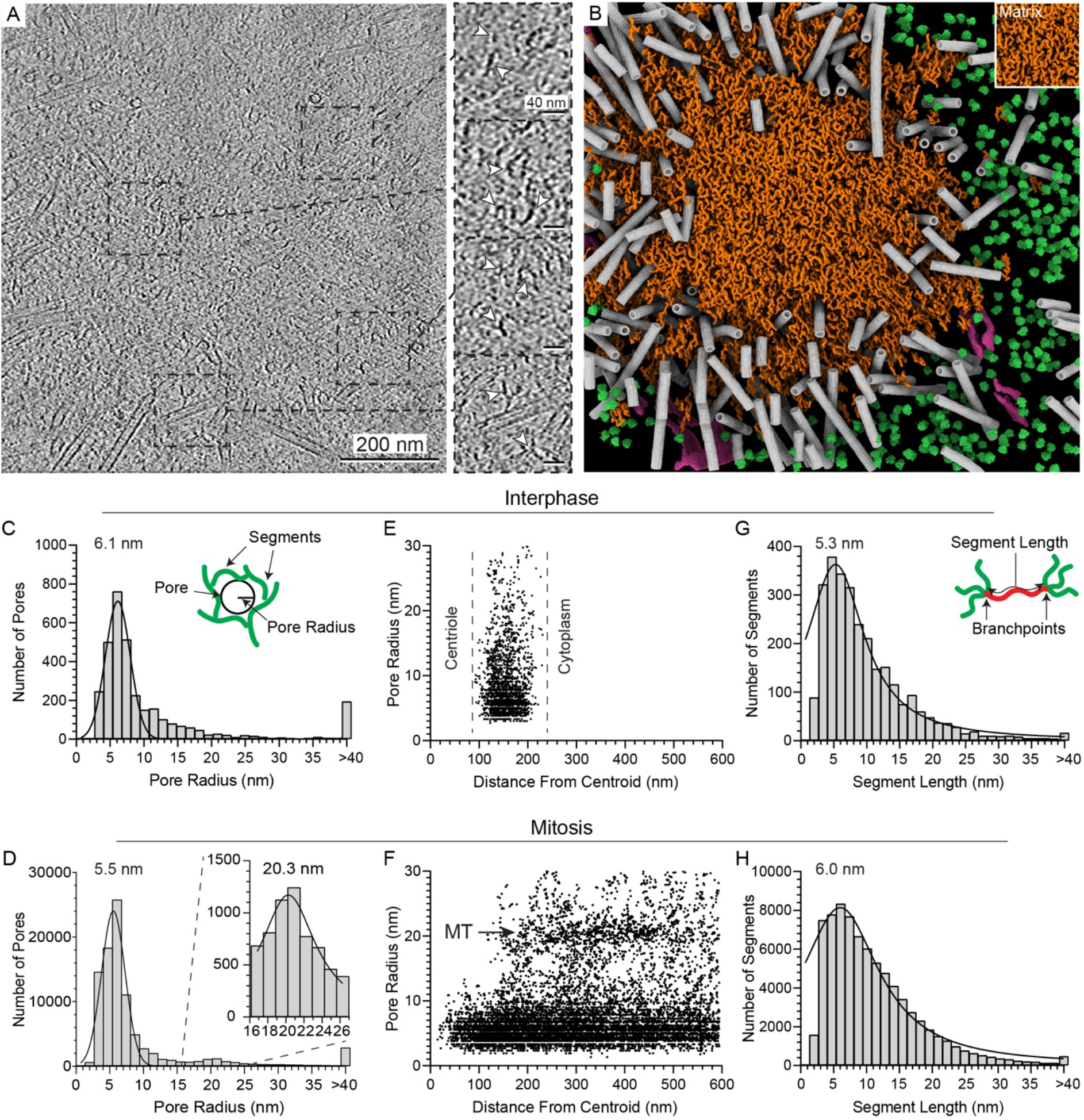
Architectural properties of the centrosome matrix. A) A denoised tomographic slice of a metaphase centrosome. Filaments highlighted in the magnified insets of the regions delineated by boxes. B) Segmented volume of a PCM-localized matrix, in the context of other cellular structure using the same color scheme as in Figure 1B. The remaining tomograms and segmentations analyzed in this work are provided in Supplementary Figure 5. C, D) Histograms of pore radii in interphase (C) and mitosis (D), with a Gaussian fit applied. Mean values are indicated. Schematic shows how pores were defined. Histogram from mitotic PCM includes a label for a secondary peak that corresponds to regions occupied by microtubules. E, F) Radius of pores along the distance from the center of the centriole (interphase: E) or ribosome-excluded centroid (mitosis: F). Gaps along the Y-axis are a result of the possible combinations of integer values in the distance matrix with finite sampling. Dashed lines in interphase correspond to the start and stop of the matrix in the data, as marked by ‘Centriole’ and ‘Cytoplasm’. MT in mitosis corresponds to the width of microtubule masking. G, H) Histograms of segment lengths in interphase (G) and mitosis (H) with a Lorentzian (Cauchy) fit applied. Peak values are indicated. Schematic shows how segments length were defined.

In both mitosis and interphase, the bulk of the filaments constituted an interconnected meshwork. In mitotic centrosomes, additional small segments of matrix, not connected to the main body, could be found beyond the edge of the ribosome excluded region (Figure 4B, Supplementary Figure 5B). To probe the properties of this matrix, we quantified the following parameters: the radii of pores, the lengths of segments between branch points, and number connecting segments at branch points.

Pore radii measured 6.1 ± 1.8 nm in interphase and 5.5 ± 1.7 nm in mitosis (Figure 4C, D, Supplementary Figure 5C). In mitotic centrosomes, a second peak of 20.2 ± 3.7 nm corresponds to the radius of the microtubule masks used, and therefore reflects an artefact of the analysis pipeline. Pore radii on the range of 5-6 nm are large enough to accommodate known PCM clients, including PLK-1 (7 by 6 nm, predictions from Alphafold2 (AF2) (Jumper et al., 2021)),, SPD-2 (9 by 5 nm, AF2 predictions), PP2A (5 by 4 nm, AF2 predictions), Aurora A Kinase (5 by 4 nm, AF2 predictions) and tubulin dimers (5 by 8 nm (Nogales et al., 1998)). The extended and disordered nature of some centrosome proteins, such as TPXL-1 and ZYG-9, makes it difficult to predict their ability to fit within this meshwork. Based on this matrix geometry, and assuming a non-dynamic assembly (as shown by FRAP of SPD-5 in metaphase PCM (Laos et al., 2015)), we expect complexes of the size of complete γ-TuRCs (25 by 21 nm in our average) to be excluded. However, given that γ-tubulin levels in the PCM are observed to increase even after the onset of anaphase (Mittasch et al., 2020), and microtubules are seen passing through the matrix, we hypothesize that the matrix must be flexible enough to permit passage of these large complexes.

We also examined the radius of these pores with respect to distance from the centroid, in both interphase and mitosis (Figure 4E, F), and found no change in pore size between the previously defined centrosome zones. In mitotic centrosomes, pores around 20-25 nm, corresponding to masked microtubules, are detected at distances larger than 150 nm, and a small fraction (6.8 % of the total pores) can be seen between these two peaks (between 11 and 19 nm). Therefore, most PCM molecules should be able to pass through the entire volume of the centrosome unimpeded.

We next examined the segment lengths between branch points in the matrix. These were found to be highly variable, with peaks of 5.3 nm and 6.0 nm in interphase and mitosis respectively, with broad fit widths of 5.3 nm and 7.2 nm (Figure 4G, H). This wide distribution indicates a lack of regular long-range order within the centrosome matrix. Similarly, the number of connecting segments at each branch point exhibited variability, with 11% of points containing more than 3 segments, and no detectable difference between interphase and mitotic PCM (Supplementary Figure 5F).

The PCM is known to become weaker and more brittle during the transition from metaphase to anaphase (Mittasch et al., 2020), but the underlying structural basis is unknown. We did not observe differences in either segment length or pore radius between metaphase and anaphase (Supplementary Figure 5D, E). This suggests that changes in PCM material properties are not caused by reorganization of matrix architecture but rather by changes in the strength of protein-protein associations that is not possible to resolve by our current methods.

In conclusion, we uncovered a network of PCM-resident filaments that comprise a disordered matrix. This matrix contains pores of sufficient size to accommodate key PCM client proteins (Figure 5) but may limit accessibility to larger complexes, such as γ-TuRC.

**Figure 5.**
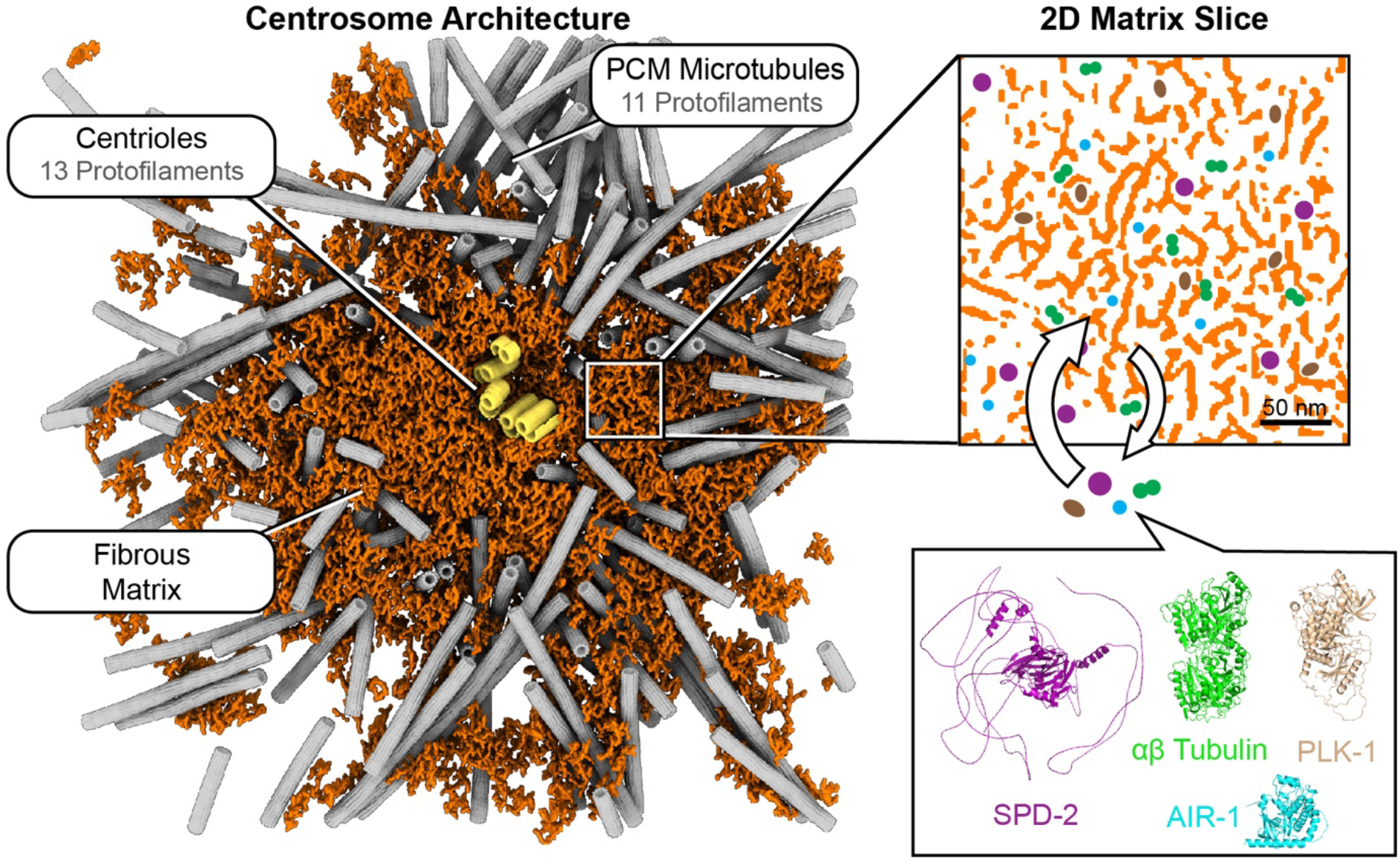
Functional properties of the PCM architecture. Left: an example of the segmented centrosome and matrix. The PCM consists of an interconnected fibrous mesh, surrounding centrioles composed of 13 protofilament microtubules. This PCM is capable of nucleating 11 protofilament microtubules by permitting access of client molecules through pores in the material (right: a 2D slice of the interconnected mesh) large enough to accommodate the client proteins SPD-2, PLK-1, Aurora A kinase (AIR-1), and tubulin dimers. Protein structures are AF2 predictions (Jumper et al., 2021), apart from the αβ tubulin structure, which is adapted from (Nogales et al., 1998).

## Discussion

Here, we present 12 *in situ* cryo-electron tomograms of centrosomes in intact *C. elegans* embryonic cells. Our data reflect a series of snapshots throughout the cell cycle and build a 3D atlas of the centrosome under native conditions. From these we observed several key structural features which help us understand how centrosomes are built and achieve function.

### Centriole maturation involves extensive structural changes

Many studies have described the structure of *C. elegans* centrioles (Dammermann et al., 2004; O’Toole et al., 2012; Pelletier et al., 2006, 2004; Sugioka et al., 2017; Woglar et al., 2022). Yet, they could not detect differences between the mother and daughter centrioles, making it unclear how centrioles mature at the structural level. Our *in situ* analysis achieved sufficient resolution with preservation of molecular detail to reveal features unique to the mother centriole: incomplete doublet microtubules, an outer ‘star’ structure, and a more pronounced central tube than the daughter. By comparison, the daughter centriole has only microtubule singlets and a prominent cartwheel, a structure likely composed of SAS-5/SAS-6, itself responsible for templating the 9-fold symmetry in the procentriole (Hatzopoulos et al., 2021; Hilbert et al., 2013; Leidel et al., 2005; Qiao et al., 2012). The cartwheel is lost in human mother centrioles (Chrétien et al., 1997), and we show the same occurs in *C. elegans*.

On the *C. elegans* mother centriole, the newly-identified partial b-tubule and outer star occupy locations which previous studies in *C. elegans* defined as the ‘paddlewheel’. Given that the paddlewheel is shown only as a fuzzy irregular density in these micrographs (Pelletier et al., 2006; Sugioka et al., 2017; Woglar et al., 2022), we suggest that it may in fact be a combination of the star and b-tubule structures blurred by fixation and/or staining artefacts inherent to room temperature EM preparations. We found the partial b-tubule to be the site of considerable heterogeneity between the different mother centrioles, with protofilament number ranging between 3 and 9. We speculate that these represent progressive maturation stages. What function does this b-tubule have within the centriole? Microtubule doublets are seen in *C. elegans* basal bodies (Nechipurenko et al., 2017) and in *Drosophila* somatic centrioles (Gottardo et al., 2015). In other organisms, a third triplet microtubule is common (Chrétien et al., 1997; Kitagawa et al., 2011; Paintrand et al., 1992). To date, *C. elegans* centrioles, commonly observed with singlet microtubules, were considered an exception. Thus, our findings support microtubule doublets as a common evolutionary structure of the centriole, conserved also in nematodes (Carvalho-Santos et al., 2011). Doublet microtubules may indeed act as the base structure of the centriole, providing it with chirality that is required for the templating of functional cilia and flagella. What is responsible for imparting microtubule doublets? The protein HYLS-1, found in embryonic centrioles, is crucial for ciliogenesis but dispensable for mitosis (Dammermann et al., 2009). Another protein that has been shown to influence doublet formation in *C. elegans* via polyglutamylation is MAPH-9, the deletion of which reduces the frequency and length of doublets in cilia. However, MAPH-9 is expressed only in ciliated sensory neurons (Tran et al., 2023), not the embryonic cells used in this study.

The molecular identity of the outer star remains unclear. Recent expansion microscopy work on *C. elegans* gonads localized many key proteins within the centriole. The outer star structure in our data localizes to the approximate positions of SAS-7, PCMD-1, SPD-5, HYLS-1, and SPD-2 (Woglar et al., 2022). Of these, SPD-5 has not been shown to be recruited specifically to mother centrioles, PCMD-1 is unstructured according to structural prediction, and HYLS-1 is required for ciliogenesis only (Dammermann et al., 2009). Thus, these proteins are unlikely to comprise the star structure. In contrast, SAS-7 has been implicated in forming part of the paddlewheel (Sugioka et al., 2017) and SPD-2 localizes most closely to the paddlewheel in the expansion microscopy study (Woglar et al., 2022). Thus, we speculate that SPD-2 or SAS-7 may form this outer star, and future studies employing the methodologies developed in our work on SPD-2 or SAS-7 conditional mutants may be able to address this question. The function of this structure also remains an open question. On the one hand, it may serve to mechanically stabilize the exceptionally small *C. elegans* centriole, substituting for A-C linkers found in higher eukaryotes (Le Guennec et al., 2020). On the other hand, it may form the platform for the tethering and assembly of the PCM on the mother centriole. The star structure may also be a primitive form of distal appendages, which have not been previously reported in nematodes.

### *C. elegans* centrosomes spatially regulate microtubule protofilament number

Purified *C. elegans* tubulin assembles into microtubules with heterogeneous geometries *in vitro*, with an average of 12.6 protofilaments and a range between 11 and 15 (Chaaban et al., 2018). Our data revealed that *C. elegans* microtubules have consistent and spatially-controlled geometries, as centriole microtubule have 13 protofilaments, whereas microtubules found in and around the PCM all had 11 protofilaments. These findings suggest the involvement of specific factors that constrain microtubule geometry *in vivo*. Here, we identified capping structures at the minus ends of PCM microtubules. The microtubule-proximal end of this cap matches the 11-fold symmetry of the microtubule. The microtubule-distal end of the cap exhibits a conical shape that resembles the GCP-containing, anchoring end of human y-TuRC, which is known to bind microtubule minus ends (Liu et al., 2020; Moritz et al., 2000; Wieczorek et al., 2020). We propose that these caps represent atypical y-TuRCs with 11-fold symmetry, which have yet to be reported in *C. elegans* or elsewhere. It is therefore likely that tubulin templating by this atypical y-TuRC establishes the consistent 11 protofilament geometry of PCM microtubules.

Conversely, both mother and daughter centriole microtubules consistently contained 13 protofilaments. How can a system assemble microtubules with different protofilament numbers within nanometer-scale proximity? γ-TuRC likely does not establish the 13 protofilament geometry of centriole microtubules in *C. elegans* embryos for two reasons: 1) we did not detect any y-TuRCs on centriole microtubules, and 2) all potential y-TuRCs that were detected were associated with 11 protofilament microtubules in the PCM. Although RNAi knockdown of γ-tubulin results in elongated centrioles, this could be a side effect of an increased pool of soluble α/β tubulin dimers caused by downregulation of PCM-based microtubule nucleation (O’Toole et al., 2012). Templating of centriole microtubules therefore may arise from another, unknown mechanism possibly involving SAS-4 (Pelletier et al., 2006). Alternatively, posttranslational modification (Cueva et al., 2012) and incorporation of β tubulin isoforms (Ti et al., 2018) could also constrain protofilament number, as demonstrated in other organisms.

### The PCM consists of a disordered porous meshwork

Prior to this study, conflicting models were proposed to describe the formation, architecture, and behavior of PCM, especially in mitosis (Raff, 2019; Woodruff, 2021). Specifically, it was unclear if the mitotic PCM comprises a regular lattice, an irregular gel-like network, or a disorganized liquid phase. Here, we show the presence of a disordered, interconnected meshwork throughout the volume of the PCM, in both interphase and mitosis. The properties of this matrix were remarkably consistent, with mean pore radius and segment length changing by only 0.6 nm and 0.8 nm, respectively, as the centrosome matures. The measured matrix properties are invariant even between metaphase and anaphase, when the PCM weakens dramatically (Mittasch et al., 2020). As it is difficult to infer the dynamic material properties of a substance from static snapshots, *in vivo* rheology is required to adequately test if the PCM transiently exhibits liquid-like or gel-like behavior at different stages of maturation.

Ultimately, it was not possible to assign molecular identity to the meshwork based on the segment lengths and branch numbers due to their variability. However, prior evidence would implicate SPD-5, as this protein uses multivalent coiled-coil interactions to assemble into micron-scale scaffolds that are capable of recruiting PCM client proteins (Rios et al., 2023; Woodruff et al., 2017, 2015). This could in theory give rise to a meshwork of variable pore sizes and segment lengths. How conserved is the architecture of the PCM? Outside of *C. elegans*, it is known that PCM scaffold proteins, such as Cnn (Timothy et al., 1999) and CDK5RAP2 (Fong et al., 2007), have limited sequence homology but exhibit similar secondary structure consisting of a series of coiled-coils connected by disordered linkers. This coiled-coil-linker arrangement could lead to similar PCM architectures. However, the differences between these scaffolds may also lead to architectural differences in pore size and segment length, potentially explaining the difference in segment length reported here in *C. elegans* (6.0 nm in mitosis) and that previously reported for *S. solidissima* (12-15 nm) (Schnackenberg et al., 1998). Thus, the underlying interconnected porous meshwork shown in this data for *C. elegans* may be reflective of a common mechanism for many species.

### Perspective

While *C. elegans* has served as a prominent model in the study of centrosome biology, as a multicellular organism, it also presents technical challenges for *in situ* cryo-ET investigations aiming at molecular resolution. Here, we deployed an alternative preparation to generate single cells from embryos and construct a 3D molecular atlas of centrosomes along the cell cycle stages. This approach, and new technological breakthroughs in the field enabling the preparation of vitrified lamellae directly from the intact frozen-hydrated organisms (Schiøtz et al., 2023), will enhance the study of key structures in other multicellular models. Our data provide a first glimpse into fine architectural features of conserved cellular structures, including the centrioles, cytoplasmic microtubules, the γ-TuRC and PCM. These data establish a reference for future comparison across evolutionary diverse organisms.

## Materials and Methods

### Worm strains, embryo collection, and embryo dissociation

*C. elegans* worms expressing GFP::SPD-5 and H2B::mCherry (JWW69: unc-119(ed9) III; ltSi202[pVV103/ pOD1021; Pspd-2::GFP::SPD-5 re-encoded;cb-unc-119(+)]II; ltIs37 [(pAA64) pie-1p::mCherry::his-58 + unc-119(+)] IV, described in (Woodruff et al., 2015)) were maintained at 22°C on Nematode Growth Media (NGM) with OP50 until reaching the adult gravid stage, as described in (Brenner, 1974).

For each preparation, at least four 100 mm petri dishes with worms were grown until the bacteria was almost fully consumed to ensure higher worm numbers while not starving the worms. Plates were washed off with M9 media (Brenner, 1974), supplemented with 0.1% polyethylene glycol (PEG) 3350 (Sigma # 88276-250G-F), and the washed material run through a 40 μm cell strainer (Falcon #352340) to remove leftover bacteria, eggs and larvae. The material retained on the strainer (gravid mothers) was then washed into 1.5 ml Eppendorf tubes and washed 3 times by centrifugation at 600 RCF for 6 min at 22°C, by aspirating the supernatant and addition of 1000 μl M9 with 0.1% PEG. Embryos were enriched through bleaching of gravid mothers. In brief, 1.8 ml of bleaching solution was freshly prepared by combining 750 μl H_2_O, 600 μl 5 M NaOH, 450 μl 5% bleach (Thermo Fischer Scientific #419550010). Worms were split such that ∼10 μl of worm pellet was visible per tube. To each tube, 450 μl M9 with 0.1% PEG and 150 μl bleach solution was added, and the tubes shaken on a benchtop thermomixer at 1400 RPM and 22°C for 13 min. The suspension was immediately transferred to a benchtop centrifuge and spun at 600 RCF for 6 min at 22°C, the supernatant removed, and embryos resuspended in M9 with 0.1% PEG. The M9 media was then replaced with L-15 media without phenol red (Thermo Fischer Scientific #21083027) supplemented with 50 U/ml pen-strep mix (Thermo Fischer Scientific #15140-122), 10% FBS (Thermo Fischer Scientific #11550356), and sucrose (Fisher Scientific #AAJ21938A1) up to 340 mOsm. This was done by centrifugation at 600 RCF for 6 min at 22°C, replacing the supernatant with modified L-15, for a total of three times. After the last centrifugation, the embryos were pooled into a single tube, centrifuged, and the bottom 200 μl retained.

The following was modified from (Christensen et al., 2002), to better suit a cryo-ET workflow. 200 μl of 4 mg/ml chitinase (Sigma Aldrich #C6137-25UN) dissolved in modified L-15 media was added to the embryo suspension and rocked for 15 min. This was followed by two rounds of centrifugation at 300 RCF, 3 min, 22°C, and replacing the supernatant with supplemented L-15. After the final centrifugation, only the bottom 50 μl was left in the Eppendorf tube. Cells were separated by gently aspirating and dispensing with a 200 μl pipette tip for 10 min. 25 μl accutase solution (Stemcell Technologies #07922) was added and left to incubate at room temperature for 10 min with gentle shaking. Cells were passed through a 200 μl pipette tip for another 10 min. To remove larger cell clumps, cells were filtered through a 20 μm and 10 μm cell strainer (pluriSelect #43-50020-03 and #43-50010-03, respectively). Cells were examined by light microscopy (20X air objective) to ensure that cell clusters were smaller than 20 μm.

For live cell light microscopy experiments, the above was carried out under a sterile cell culture hood.

### Live cell light microscopy

Prior to embryo dissociation, 35 mm low μ-Dish with a polymer coverslip (Ibidi #80136) were prepared by acid treatment overnight in 1 M HCl at 50°C, washed with dH_2_O, left to dry for 1 hour under a sterile cell culture hood, and each incubated with 300 μl 0.1% poly-l-lysine (Sigma #P8920) for 1h. Dishes were washed 5 times with 300 μl dH_2_O. Dissociated cells were pipetted into the treated dishes, and the total volume of 200 μl supplemented by addition of modified L-15 media. Cells were allowed to settle for 1h, and washed in 200 μl fresh media 5 times to remove unattached cells.

Imaging was carried out on a Zeiss 780 confocal microscope with a plan-apochromat 63x oil immersion DIC objective lens, NA 1.2, with the chamber set to 21°C. Regions containing cell clumps were imaged under the following conditions: pixel spacing of 0.264 μm, image size 512 x 512 pixels (134.95 μm x 134.95 μm), pixel dwell time of 3.14 μs, pinhole set to 288 μm, with 8 X 0.5 μm Z steps. Power of the 561 nm laser was set to 2.5%. Sites were imaged at 5-min intervals for 100 min.

### Plunge freezing

Plunge freezing was carried out within an hour from cell dissociation, to ensure cells were viable. Quantifoil Holey SiO_2_ R 1/4 Au 200 mesh EM grids (Quantifoil Micro Tools #N1-s13nAi20-01) were glow discharged on both sides in a Pelco easyglow for 45 s at 0.37 mBar and 15 mA. Plunge freezing was carried out using a Leica EM GP2. Chamber temperature was set to 22 °C and humidity to 70%. 4 μl of cell suspension was added to the front of the grid, and 2 μl of supplemented L-15 media to the back side to aid blotting. Grids were blotted from the back for 3 s and plunged into liquid ethane at - 185 °C. Grids were stored in grid boxes in liquid nitrogen until further use.

### Cryo-fluorescence microscopy

Grids were clipped into Autogrid cartridges modified for shallow angle FIB milling, mounted on a dedicated shuttle and transferred into a Leica cryo-confocal microscope based on the Leica TCS SP8 system, equipped with a cryo-stage. Imaging was performed with a 50x NA 0.9 air objective. Grid overviews were acquired in epifluorescence mode and montaged (Mosaic stitching module in Leica Application Suite X 3.5.5.19976) to generate a map from which regions of interest were identified. These regions were imaged in confocal mode with the following conditions: image size 1056 x 1056 pixels (182.65 μm x 182.65 μm), with 4x line integration, and mirror set to speed 200 Hz giving a pixel dwell time of 2.03 μs. The pinhole was set to 91.95 μm with 55 0.37 μm z steps, crossing 20.11 μm total. Both the 488 nm and 552 nm lasers were set to 50% power, with detection from 492 to 540 nm and 596 to 719 nm. Grids were stored in liquid nitrogen until further use.

### Cryo-focused ion beam milling

Cryo-FIB milling was carried out on a dual beam Aquilos1 equipped with a cryo-transfer system and stage (ThermoFisher Scientific). Prior to milling, an overview scanning electron microscopy image of the grid was manually overlaid with the montage from the cryo-fluorescence microscopy, and grid squares containing cells of interest defined. Confocal microscopy stacks were projected in Fiji 2.9.0 (Schindelin et al., 2012), and the Z projection manually overlaid to the electron beam images.

Lamellae milling positions were recorded using Maps 2.5 software (Thermo Fischer Scientific). Grids were coated with organometallic platinum using the Gas Injection System for 10 sec, and sputtered with metallic platinum at 10 Pa, 1 kV, and 10 mA for 20 s. Milling was carried out at a 20° angle stage tilt using a shuttle with a 45° pretilt at 30kV. 300 nm wide micro-expansion joints were milled on both sides of the target lamella to reduce the effects of stress (Wolff et al., 2019). Milling was performed at currents of 1 nA, then 500 pA and 300 pA at a distance of 5 μm, 3 μm and 1 μm from the position of the target lamella, respectively, ablating material above and below the region of interest. Fine milling was done at a current of 100 pA, milling first below the lamella and then above. Lamella were milled either fully manually or with automated rough milling using SerialFIB (Klumpe et al., 2021). Grids with lamellae were sputter coated for 3 s at 10 Pa, 1 kV, and 10 mA.

### Cryo-ET data collection

Data collection was carried out on a Titan Krios (ThermoFisher Scientific) cryo-TEM at 300 kV, equipped with a field-emission gun, a Gatan quantum post-column energy filter operated in zero-loss, a K2 Summit direct electron detector (Gatan), and a Volta phase plate (ThermoFisher Scientific). Grid maps at 233 nm/pixel and lamella maps at 2.2 nm/pixel were first collected. These were overlaid with the correlative cryo-light microscopy montage and the projected confocal stack, respectively, to aid target tilt-series acquisition.

SerialEM (Mastronarde, 2018) was used to collect dose-fractionated tilt-series images, using a tilt range of −60° to +60° at 2° increments with the sample pre-tilted to compensate for the FIB milling angle, in a dose symmetric scheme (Hagen et al., 2017). A pixel size of 3.37 Å/pixel, a defocus range between 2.5 μm and 4.5 μm, and a total electron dose of 120 e^-^/ Å^2^ were applied. In total, 12 tomograms of centrosomes acquired with a Volta phase plate preconditioned for 6 min were analyzed in this work.

### Tomogram reconstruction and denoising

Motion correction and CTF estimation were carried out in Warp 1.0.9 (Tegunov and Cramer, 2019), starting from processing original frames. Tilt-series alignment of stacks written in Warp was carried out using etomo (Kremer et al., 1996). Final tomogram reconstruction was carried out in Warp, with a voxel size of 13.4 Å/pixel (4x binning) to enhance the signal-to-noise ratio and reduce overall tomogram size. For segmentation of the centrosome matrix, we employed denoising using cryoCARE 0.2.1 (Buchholz et al., 2018). For each tomogram to be denoised, two tomograms were reconstructed using odd and even frames respectively. These tomograms were also deconvolved in Warp, and a lamella mask generated using TOM toolbox in Matlab (Nickell et al., 2005) was applied. For tomograms with stronger large intensity gradients due to curtaining effects on the lamella, filtering with kernel size 10 was applied to the denoised tomograms using TOM toolbox, and this filtered tomogram (background) was subtracted from the original tomogram.

### Segmentation of microtubules, centrioles, ribosomes, and membranes

Segmentation of microtubules in 11 tomograms was carried out using a cylinder correlation function in Amira 2021.1 (Stalling et al., 2005; Rigort et al., 2012; Weber et al., 2012). Raw 4x binned tomograms were Gaussian filtered with a kernel of 3 voxels prior to segmentation. Cylinder correlation was run using a cylinder length of 53.92 nm (40 voxels), angular sampling of 8 degrees, mask cylinder radius of 13.48 nm (10 voxels), outer cylinder radius of 12.13 nm (9 voxels), and inner cylinder radius of 6.7404 nm (5 voxels). Missing wedge correction was set in the Y-tilt axis from −60° to +60°. Correlation tracing had a minimum seed correlation of 60, minimum continuation quality of 60, direction coefficient of 0.3, minimum distance of 24.27 nm (18 voxels), and minimum length of 53.92 nm (40 voxels). The search cone was 53.92 nm (40 voxels) long, 37° wide, with a minimum step size of 10%. Results were manually examined in the filament tab, and erroneous results were removed. To smooth the segmentation results and connect breaks in the microtubules, this output was run again through cylinder correlation: a surface for the first round of cylinder correlation was extracted with tube scale 8. This surface was placed into a volume with dimensions corresponding to the tomogram, and subjected to cylinder correlation with the same parameters as above, but with the inner cylinder radius set to zero. For the segmentation of the remaining tomogram, binary masks corresponding to the output of the first 11 were used to train a DeePiCt model for microtubules (https://github.com/ZauggGroup/DeePiCt) (de Teresa-Trueba et al., 2023), and the trained model (11cents5inter6mitotic_pp, bundled with DeePiCt) used for prediction on the remaining tomogram. The outputs were visually inspected and cleaned in Amira to remove false positives.

Centriole microtubules were manually traced in Amira 2021.1.

Ribosomes and membranes were segmented in DeePiCt, using the models ribo_model2_IF4_D2_0 and full_vpp_memb_model_IF4_D2_BN respectively (https://github.com/ZauggGroup/DeePiCt). Results were inspected in Amira to remove false positives.

### Analysis of centrosomal zones

Analysis was conducted in MATLAB 2019a. For mitotic centrosomes, centroids for the ribosome and microtubule exclusion regions were calculated as follows: binary segmentations for each of the structures were averaged along the Z-axis to obtain a 2D projection. The diameter and approximate centroid location of the exclusion region were estimated from this projection. A circle with a diameter approximately 50% larger than the estimated excluion region was drawn from a putative centroid. The segmentation within this circular mask was inverted, assigning a value of zero to the signal and one to the background. The centroid coordinates in X and Y of the inverted segmentation was calculated iteratively until convergence. To determine the centroid value in Z, a single round of the above process was performed with X and Y values fixed to the 2D output. For interphase centrosomes, centroids were calculated from the centroid of a binary centriole segmentation, from which spheres were grown concentrically. Once a sphere encountered density from the binary segmentation for each of the structures, the sphere’s radius and the number of voxels were recorded.

### Subtomogram analysis of microtubules

Coordinates and initial Euler angles of particles were assigned based on the microtubule segmentations using a custom script in MATLAB 2019a (https://github.com/ZauggGroup/DeePiCt) (de Teresa-Trueba et al., 2023). The outputs of the scripts provide both a table file for use in dynamo1.1.514 (Castaño-Díez et al., 2012), and a STAR file for use in Warp 1.0.9 (Tegunov and Cramer, 2019). The table includes unique identifiers for each tomogram, and each microtubule. 19,339 subtomograms were reconstructed by sampling at every 8.09 nm along the traced microtubules in Warp using a pixel size of 6.74 Å and a box size of 72 voxels. The subtomograms were then imported into dynamo via the dynamo GUI and the third Euler angle randomized prior to alignment.

Subtomograms were aligned in dynamo across 2 rounds of 5 iterations each. The first round aligned subtomograms around the new Z, with a 30° search cone, but no azimuthal searches. The second round searched only around the azimuth, with a 18° (360/11) search range. Both rounds allowed 53.92 Å (8 voxel) shifts from the center of the first iteration. The resolution of the resulting average was determined to be 26 Å from maps reconstructed from odd and even half-sets using a MATLAB script with an FSC cutoff of 0.5 (half-sets were not processed independently).

Next, per-microtubule averages were generated using the unique microtubule identifiers in the table. To improve contrast, these per-microtubule averages were projected along the Z axis to create a 72×72 pixel 2D average. Protofilament number was determined in MATLAB 2019a, by comparing each average to reference densities of microtubules with 11 to 15 protofilaments (EMD-5191, EMD-5192, EMD-5193, EMD-5194, EMD-5195 (Sui and Downing, 2010)). These reference EM maps were resampled to 6.7404 Å/pixel using RELION image handler and averaged along their long axis to obtain 2D images. Each microtubule average from our data was rotated until it returned the highest cross-correlation coefficient for each reference. The highest cross-correlation obtained from matching against the different structures was recorded as the protofilament number for the microtubule. Similarly, microtubule polarity was determined by comparison to a microtubule reference with positive polarity (minus side at the centrosome and pointing away) by taking the 2D projection of the longest single microtubule seen in the data. This reference was flipped along its Y axis to serve as the reference for negative polarity. All projected microtubule averages were rotated against both templates until they recorded the highest cross correlation, and the highest score recorded as the polarity for each microtubule.

### Centriole analysis

Centrioles were circularized and symmetrized in Fiji (Schindelin et al., 2012) using CentrioleJ (Guichard et al., 2013). To determine the lengths of different structural features through the centriole, the tomograms were viewed in 3dmod (Kremer et al., 1996) and positioned in the slicer window such that the feature of interest along each microtubule lay flat and fully visible. Line scans of the generated images were then produced in Fiji, and the length of each feature was determined as the distance in the scan where the mean pixel value is lower (higher density) than the mean pixel value of the entire image.

### Centriole subtomogram averaging

Similar to the microtubule subtomogram averaging, coordinates and initial Euler angles of particles were assigned from the manual segmentations by use of a custom script in MATLAB 2019a outputting both a STAR and table file. 800 subtomograms were reconstructed from 12 centrioles every 8.09 nm in Warp 1.0.9 with pixel size of 6.7404 Å and a box size of 180 voxels. Unique identifiers for the tomogram number, centriole number, and microtubule number within the centriole were recorded in the table. The third Euler angle for each centriole microtubule was initially fixed relative to the other microtubules in the same centriole. To align these microtubules to one another, the third Euler angle for each sutomograms was rotated 40° (360°/9) relative to the microtubule before it. Masking and alignment were focused on the a-tubule. An alignment job was then run in dynamo 1.1.514, 12 iterations allowing shifts of 40.44 Å (6 voxels), with cone and azimuth search ranges of 27.7°. The generated average was then split into different centrioles using the table identifiers, and then projected along Z to improve contrast for analysis of the visualized features. The resolution of the resulting global average was determined to be 44 Å using a MATLAB script, using an FSC cutoff of 0.5 (odd and even half-sets were not processed independently).

### γ-TuRC subtomogram averaging

281 particles were manually picked at ends of short microtubule segmentsfrom 4x binned denoised tomograms. Subtomograms and 3D CTF models were reconstructed in Warp 1.0.9 with voxel size of 13.48 Å and 6.74 Å (4x binning and 2x binning, respectively) with box sizes of 74 and 148 voxels, respectively. Initial angles were determined by alignments in dynamo 1.1. In RELION 4.0.1 (Kimanius et al., 2021), 4x binned subtomograms were first subject to a 2D classification job, which resulted in all 281 particles being assigned to a single class. The alignment results were then refined at 2x binning, with a 450 Å spherical mask over the tip and 15° angle restriction, to obtain the final 38 Å resolution average. Resolution was determined using RELION postprocessing, using a ‘gold standard’ FSC cutoff of 0.143.

γ-TuRC orientations were determined in MATLAB by determining the vector between the ribosome excluded centroid to each γ-TuRC in the respective volume, and the vector along the long axis of each γ-TuRC (as determined from the Euler angles). Angles were determined by taking the dot product between these and dividing it by the product of the magnitudes of each vector.

### Subtomogram Averaging of Ribosomes

Ribosomes were aligned and averaged solely for improving the quality of depictions presented in this manuscript. Using the coordinates for ribosomes as identified as described in the segmentation section, 10,220 subtomograms and 3D CTF models were reconstructed in Warp 1.0.9 at 10 Å/pixel and a box size of 44 voxels. Subtomograms were subject to 3D refinement in RELION 4.0.1 (Kimanius et al., 2021), with no mask using the default parameters.

### PCM Matrix analysis

Denoised tomograms were used for matrix segmentation. Centrioles, microtubules, ribosomes and membranes were all masked in the tomogram volumes using MATLAB 2019a by expanding the segmentation of these features by 24, 11, 8, and 4 voxels respectively. The expansion was necessary to ensure all densities corresponding to each feature have been removed, as missing wedge artefacts can cause elongation along the tomogram Z direction. Additionally, strict lamella masks were applied. Two models for the matrix were generated following manual annotation using Ilastik 1.3.3. One utilized 3D Pixel Classification with selected features and 7 labels, while the other used 2D Autocontext trained on the central 50 XY slices of the tomogram. The outputs of the Autocontext models, which were determined to provide a better representation of the meshwork following visual inspection, were combined, binarized in MATLAB 2019a to obtain the final matrix model. For mitotic centrosomes, we applied an additional spherical mask around the centrosome to remove excess signal from outside the centrosome. The γ-TuRC averages were placed into the matrix model and then subtracted from it.

Pore sizes were determined by running a distance transformation of this matrix, and identifying local maxima in MATLAB 2019a. Radii of the pores were calculated by multiplying the distance transformation by the regional maxima. A Gaussian fit of the data was applied using Graphpad Prism 9.5.1.

For determining segment lengths, the matrix was skeletonized using the Skeleton3D function in MATLAB 2019a. Segment lengths were calculated using the Skan toolbox (0.11.0) (Nunez-Iglesias et al., 2018) in Python 3.10.9, considering only segments between junctions. Connections shorter than 2 voxels were removed from the data. Fitting was done in Graphpad Prism 9.5.1, using a Lorentzian fit, with the constraints that the amplitude, center, and width are larger than one. The overflow bin was not included in this fitting.

## Supplemental material

Supplemental material associated with this manuscript includes Supplementary Figures 1-5.

## Data availability

Subtomogram averages generated in this work are deposited to the Electron Microscopy Data Bank (EMDB) with the following accession numbers: centrioles EMD-19779, microtubules EMD-19778 and γ-TuRC EMD-19780. A representative tomogram is deposited on EMDB with accession number EMD-19781. The cryo-ET data presented in this work, including raw frames, tilt-series, reconstructed tomograms and corresponding annotations, will be deposited on the Electron Microscopy Public Image Archive (EMPIAR). All entries will be released upon publication. Cryo-EM maps of microtubules with 11 to 15 protofilaments were obtained from EMD-5191, EMD-5192, EMD-5193, EMD-5194, EMD-5195. The atomic model of the human γ-TuRC was obtained from the PDB 6V6S.

## Acknowledgements

We thank the Mahamid group members for fruitful discussions and help, especially Marie Spindler and Rasmus Jensen for providing valuable input on the manuscript and analysis, and Frosina Stojanovska for assistance with python scripts. We thank the EMBL cryo-EM platform, in particular, Wim Hagen, for support in cryo-EM data acquisition, the EMBL Advanced Light Microscopy Facility for support in fluorescence image acquisition and analysis, Thomas Hoffmann and EMBL IT for computational support, Simone Koehler and Ana Rita Rodriguez Nevez for help with *C. elegans* maintenance. The cryo-confocal microscope was developed in collaboration with Leica Microsystems, and we thank Jan De Bock and Martin Schorb for invaluable support. We thanks Niccolo Banterle for critical input on the manuscript. J.B.W. is supported by a Cancer Prevention Research Institute of Texas (CPRIT) grant (RR170063), a Welch Foundation Grant (I-2052-20200401), an R35 grant from the National Institute of General Medical Sciences (1R35GM142522), and the Endowed Scholars program at UT Southwestern. M.U.R. was supported by a National Research Service Award T32 (GM007062). J.M. was supported by a European Research Council starting grant to (3DCellPhase^-^ 760067) and the EMBL.

## Author contributions

F.T., J.B.W., and J.M. conceived this study. F.T., J.B.W., and M.U.R. optimized the *C. elegans* embryo dissociation and cell culture. F.T. performed all sample preparation, data acquisition, and analysis. E.Z. performed live cell imaging. F.T., J.B.W., and J.M. wrote the manuscript with input from all co-authors.

