## Supplementary Figures 1-5 for "Native molecular architectures of centrosomes in *C. elegans* embryos"

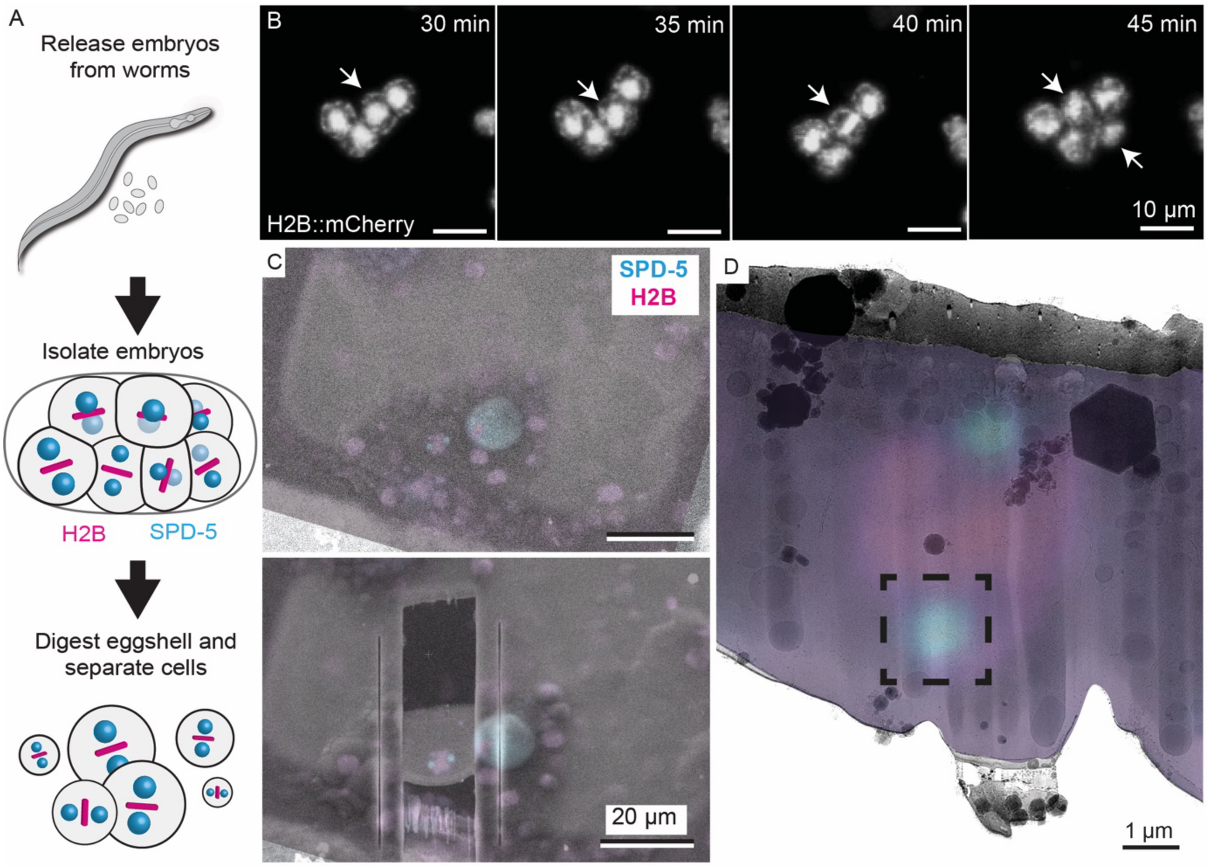


Supplementary Figure 1. CLEM-guided sample preparation and data collection. A) Schematics of the embryonic cell dissociation approach. B) Live cell fluorescence light microscopy of a cell cluster obtained following the pipeline shown in A. Arrows indicate a cell undergoing mitosis. Indicated time points are from the start of data acquisition. C) Cryo-scanning electron microscopy (SEM) images showing a cell before and after FIB lamella milling. Correlated fluorescence images (maximum intensity projection of the cryo-confocal volume) with GFP::SPD-5 in cyan and H2B::mCherry in magenta are overlaid to the SEM images. D) cryo-transmission electron microscopy image of a cryo-FIB-lamella (different from that shown in C), overlaid with the correlated fluorescence data (colors as indicated in C). The dashed square indicates the location of a centrosome suitable for tilt-series acquisition and the approximate field of view of the tomographic data.

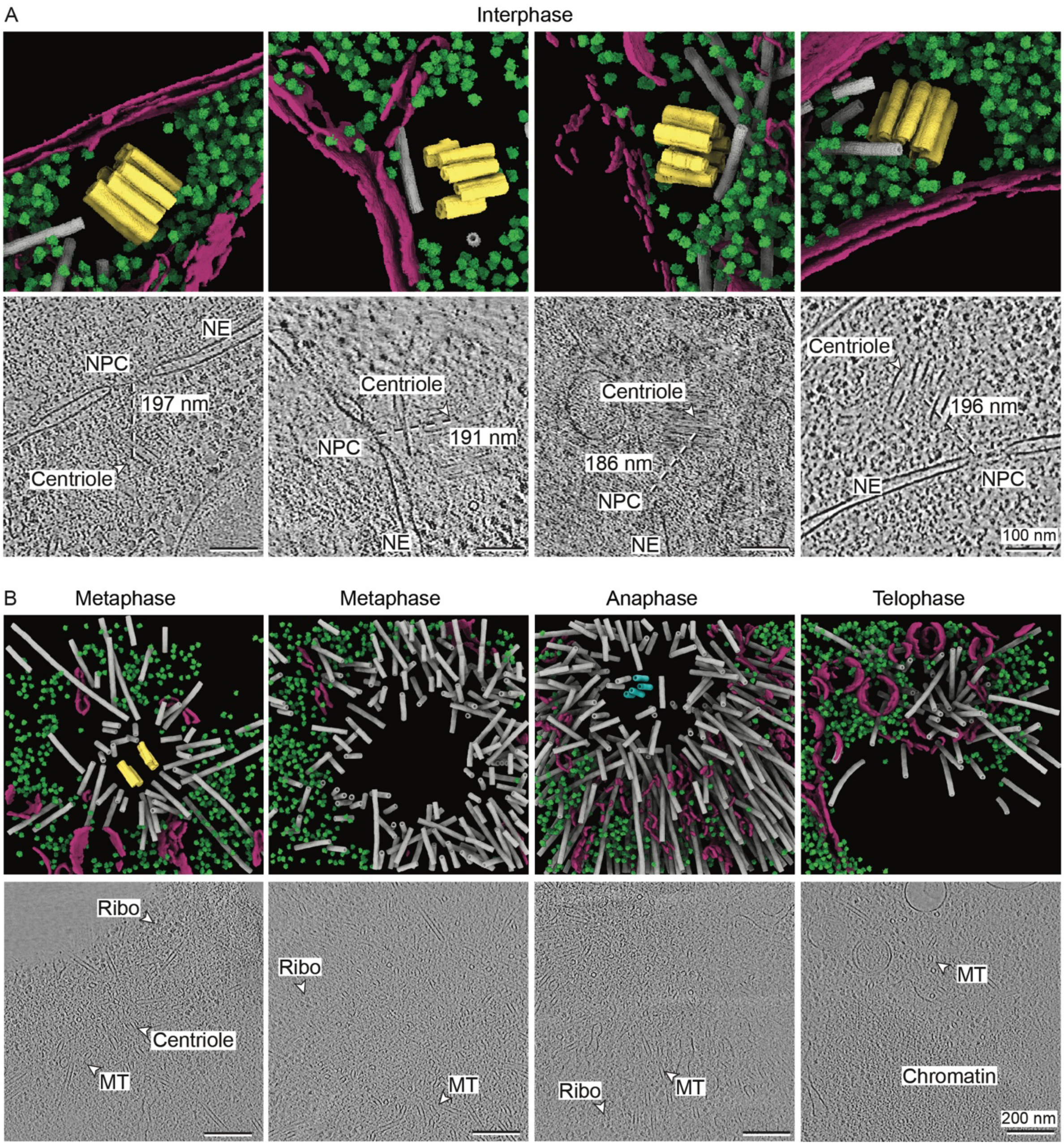


Supplementary Figure 2. A gallery of the additional centrosomes data collected in this study (complementing Figure 1). Top rows: 3D annotations: Mother centrioles: yellow, daughter centrioles: cyan, microtubules: grey, ribosomes: green, and membranes: purple. Bottom rows: tomographic slices. A) Interphase centrosomes. In tomographic slices, the Nuclear Pore Complex (NPC), Nuclear Envelope (NE), and centrioles are labeled. The dashed lines indicate the distance (186-197 nm) between the centriole centroid and NPC, with a mean of 191.5 $\pm$ 5 nm across the 5 interphase tomograms. B) Mitotic centrosomes. Cell cycle stage is indicated above the images. In tomographic slices, ribosomes (Ribo), microtubules (MT), centrioles and chromatin are labelled.

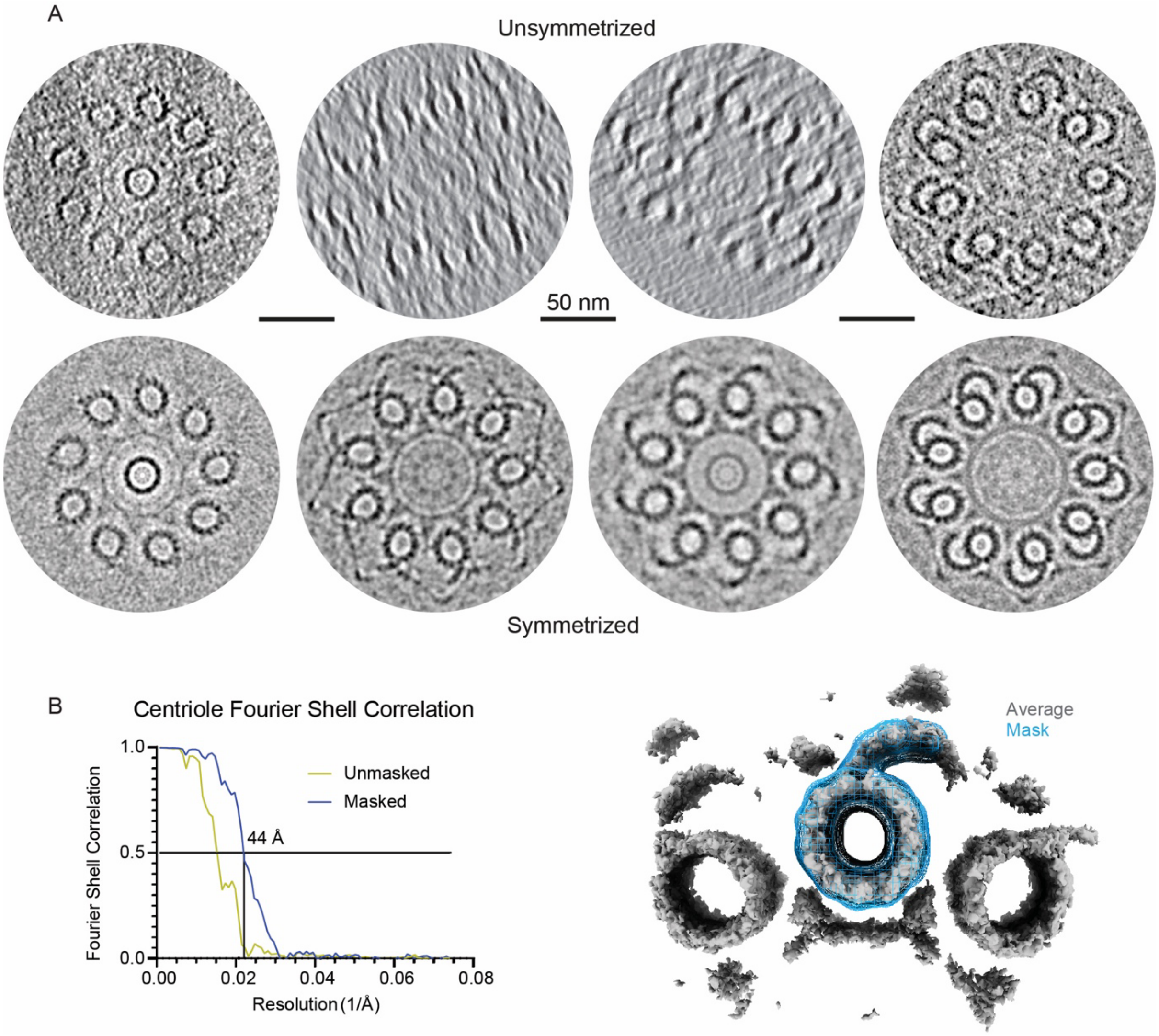


Supplementary Figure 3. Structural analysis of centrioles. A) Representation of all complete, or near complete, centrioles (i.e. not cut off by the lamella edge), imaged in this study (complementing data shown in Figure 2). Centrioles ordered from left to right by the completeness of the B-tubule. Top: raw data. Bottom: 9-fold symmetrized. B) Fourier Shell Correlation (FSC) and subtomogram average of pooled centriole data (800 subtomograms from 12 centrioles). Left: FSC curves for centriole subtomogram average shown for unmasked and masked averages, with FSC cutoff taken at 0.5 as the two half-sets were not processed independently throughout the processing pipeline. Right: The average (grey) overlaid with the mask used (blue mesh).

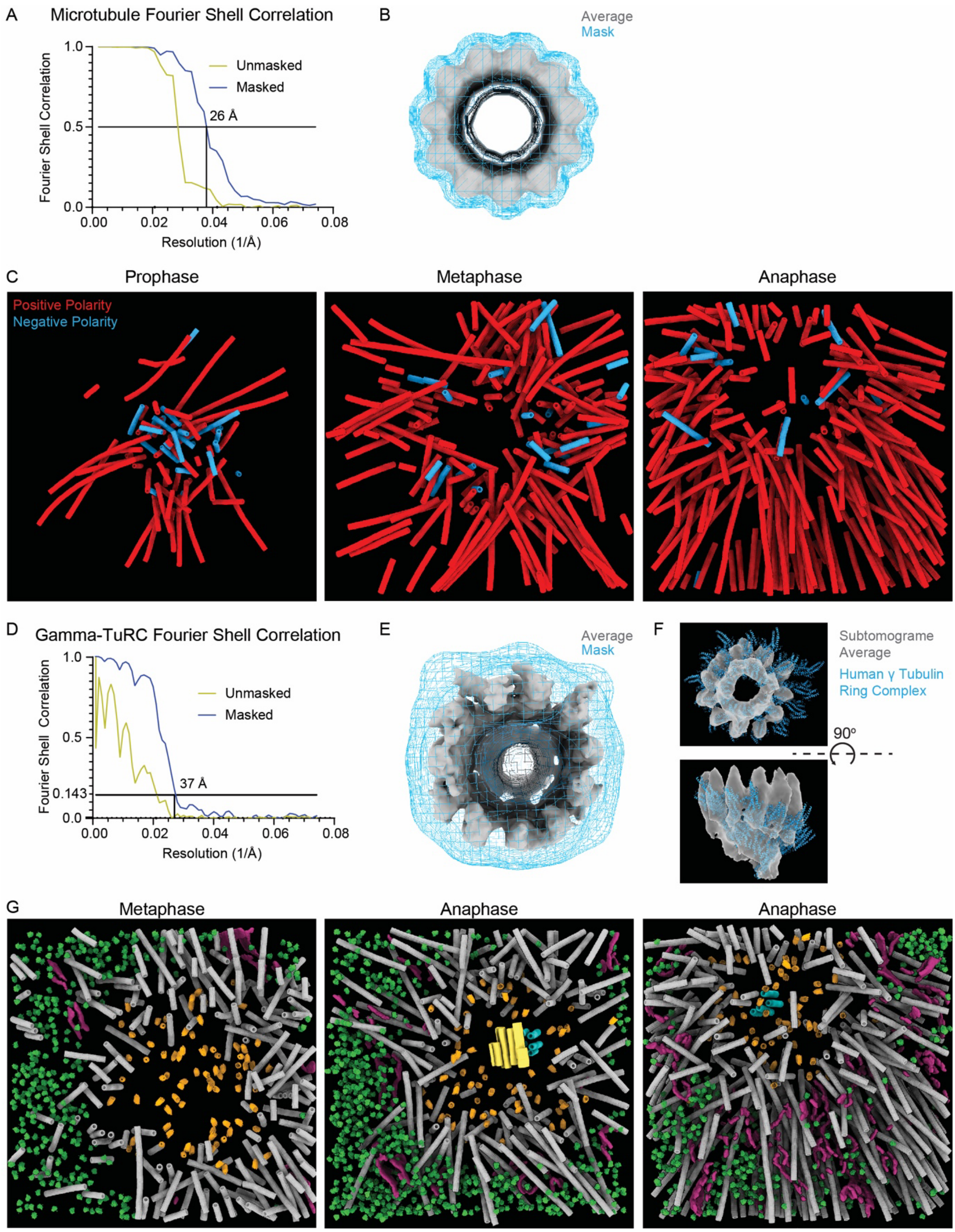


Supplementary Figure 4. Analysis of centrosomal microtubules organization. A) FSC curves for microtubule subtomogram average shown for unmasked and masked average, with FSC cutoff at 0.5 used as the two half-sets were not processed independently throughout the processing pipeline. B) The centrosomal microtubules pooled average (grey; 19,339 subtomograms) overlaid with the mask used (blue mesh). C) Examples of microtubules organization in mitotic centrosomes (cell cycles stage indicated), colored according to polarity, with blue indicating negative polarity (minus end faces away from the centroid) and red indicating positive polarity (minus end faces towards the centroid). D) FSC curve for γ-TuRC subtomogram average, with gold-standard FSC cutoff at 0.143 providing a resolution of 38 Å. Curves shown for unmasked and masked average on the right. E) The γ-TuRC subtomogram average (grey; 281 subtomograms) overlaid with the mask (blue mesh). F) A rigid body fit of the human γ-TuRC (blue; PDB 6V6S (Wieczorek et al., 2020)) into the *C. elegans* γ-TuRC subtomogram average (grey). G) Additional examples of 3D annotation of γ-TuRC complexes in mitotic centrosomes (stage indicated), complementing data shown in Figure 3. Note that γ-TuRCs were only processed and shown at ends of short microtubules.

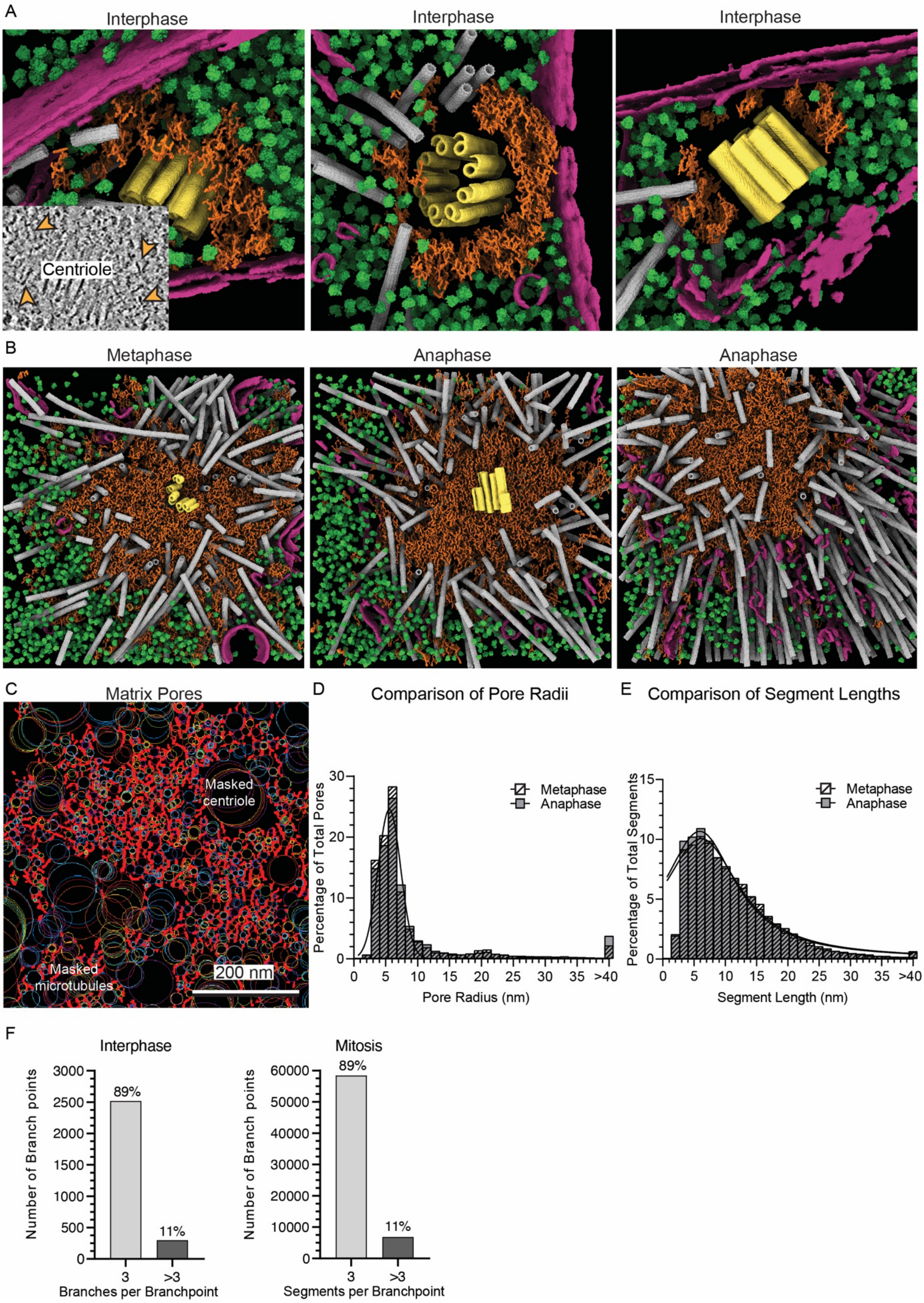


Supplementary Figure 5. Analysis of the centrosome matrix. A) 3D annotation of the PCM within interphase centrosomes analyzed in this work. Inset in left panel shows a region around the centriole with putative PCM filaments highlighted with orange arrows. Not shown on the same scale as the annotation. B) Additional examples of 3D annotation of the PCM within mitotic centrosomes analyzed in this work (complementing data shown in Figure 4). Mitotic stage is indicated above each image. C) Visual representation of the matrix segmentation and fitting for pore radius measurements. D, E) Comparison between metaphase and anaphase matrix parameters. Histograms for pore radius (D) and segment length (E) are shown with metaphase values in dashed lines and anaphase in grey. Fits for each are provided on the same graph. F) Number of connecting segments per branchpoints in the skeletonized matrix, shown for both the interphase and mitotic PCM.
